# NKCC1 as a signaling hub regulating KCC2 stability, chloride homeostasis, and seizure susceptibility

**DOI:** 10.64898/2025.12.05.692569

**Authors:** Erwan Pol, Celia Delhaye, Silvia Cases-Cunillera, Marion Russeau, Simon Blachier, Adrien Bouchet, Christophe Piesse, Johan Pallud, Gilles Huberfeld, Nicolas Pietrancosta, Sabine Lévi

**Affiliations:** ESPCI, CNRS UMR 8249, PSL Université, 10 rue Vauquelin, 75005 Paris, France; INSERM UMR-S 1270, Sorbonne Université, Institut du Fer à Moulin, 75005, Paris, France; Sorbonne Université, CNRS, Inserm, Center of Neuroscience Neuro-SU, 75005 Paris, France; Sorbonne Université, CNRS, Inserm, Institut de Biologie Paris-Seine, IBPS, 75005 Paris, France; Université Paris Cité, Institute of Psychiatry and Neuroscience of Paris (IPNP), INSERM U1266, Neuronal and Astroglial Signaling in Epilepsy and Glioma, 75014 Paris, France; Department of Neurosurgery, GHU Paris Psychiatry and Neurosciences, Sainte-Anne Hospital, 75014 Paris, France; Institute of Psychiatry and Neuroscience of Paris, University Paris Cité, INSERM U1266, IMABRAIN, 75014 Paris, France; Université Paris Cité, 75005 Paris, France; Department of Neurology, Hopital Fondation Adolphe de Rothschild, 75019 Paris, France; Sorbonne Université, École Normale Supérieure, Université PSL, CNRS, Chimie Physique et Chimie du Vivant (CPCV), 75005, Paris, France

## Abstract

Chloride homeostasis relies on the dynamic balance between the neuronal co-transporters NKCC1 and KCC2. We reveal an unexpected mechanism by which NKCC1 governs KCC2 membrane stability. NKCC1 clusters recruits SPAK and PP1 to dynamically trap KCC2, compensating for its lack of a direct SPAK-binding site. Single-particle tracking shows that these NKCC1-rich assemblies operate as signaling hubs, enabling either SPAK-driven KCC2 phosphorylation and its membrane destabilization or PP1-mediated dephosphorylation of SPAK and KCC2 membrane stabilization. Peptides that activate SPAK by engaging NKCC1’s PP1-binding motif lower KCC2 surface levels and reduce chloride extrusion, whereas a SPAK-inhibiting peptide prevents SPAK recruitment to NKCC1, stabilizes KCC2 in membrane clusters, and enhances chloride extrusion. An optimized peptide analog preserves KCC2 clustering under hyperexcitable conditions, reduces seizure frequency and severity in PTZ-induced epilepsy, and suppresses ictal activity in human epileptic tissue. These findings identify NKCC1-KCC2 coupling as a central regulatory axis for inhibitory signaling, and position our peptides as promising therapeutic candidates to restore chloride homeostasis in epilepsy and other disorders marked by impaired KCC2 membrane stability.

## Introduction

GABA type A receptors (GABA_A_Rs) mediate fast synaptic inhibition through chloride-permeable channels. The direction and efficacy of GABAergic signaling are governed by the intracellular chloride concentration ([Cl^-^]_i_), which is tightly regulated by the opposing actions of the K⁺-Cl⁻ exporter KCC2 and the Na⁺-K⁺-Cl⁻ importer, NKCC1. In epilepsies, [Cl^-^]_i_ is frequently elevated in a subset of neurons, leading to a depolarizing shift in the GABA reversal potential (E_GABA_) and impaired inhibition ^1,2^. These changes are associated with reduced KCC2 function and/or NKCC1 upregulation, and have been documented in both human temporal lobe epilepsy, gliomas, cortical malformations and relevant animal models ^3–9^. However, elevated KCC2 surface expression has also been reported in human intractable temporal lobe epilepsy^10^ and in inflammatory pain conditions such as acute arthritis ^11^. Despite the critical role of chloride homeostasis in controlling inhibition, current antiepileptic treatments primarily target neuronal excitability or GABA_A_Rs, but not chloride transporters. More than 30% of patients remain resistant to available therapies ^1^, underscoring the need for alternative strategies - particularly those aimed at restoring Cl⁻ balance. This approach could also be beneficial for other disorders involving disrupted inhibition, including neuropathies and neuropsychiatric conditions ^2^.

KCC2 membrane stability is dynamically regulated by activity-dependent signaling pathways that control its phosphorylation, lateral diffusion, and endocytosis ^12–15^. NKCC1 is likewise subject to rapid modulation via a “diffusion-trap” mechanism that adjusts its membrane clustering in response to GABAergic input ^16,17^. Both transporters are substrates of the chloride-sensitive WNK1-SPAK/OSR1 kinase cascade, which concurrently inhibits KCC2 by phosphorylating threonine Thr906 and Thr1007, and activates NKCC1 via phosphorylation of Thr203, Thr207, and Thr212, thus impairing Cl⁻ extrusion and enhancing Cl⁻ uptake ^15,18,19^. Notably, this pathway is aberrantly activated in epilepsy ^15,20^, suggesting that it contributes to pathological network hyperexcitability. A promising therapeutic strategy is to stabilize KCC2 at the membrane by inhibiting the WNK-SPAK signaling cascade. However, under conditions of elevated KCC2 surface expression ^10,11^, activation of WNK or SPAK - by promoting KCC2 membrane destabilization - could represent a novel therapeutic strategy for these diseases.

While NKCC1 contains two SPAK-binding motifs, KCC2b-the predominant neuronal isoform-lacks recognizable SPAK interaction domains ^18,19^. NKCC1 also interacts with the phosphatase PP1, which counteracts SPAK activity ^21^. We hypothesized that NKCC1 may serve as a scaffold for SPAK and PP1, positioning them in proximity to KCC2 to regulate its phosphorylation and membrane retention. Supporting this, more than 60% of NKCC1 clusters colocalize with KCC2 in cultured hippocampal neurons, forming denser and larger membrane domains than either transporter alone ^16^.

Here, we examined the behavior of KCC2 in relation to NKCC1 membrane clusters. We observed that KCC2 transporters freely diffusing in the plasma membrane become slowed and confined upon encountering regions where NKCC1 forms aggregates, suggesting that they are recruited and stabilized at these sites. Moreover, using a structure-guided design strategy, we generated membrane-permeant peptides that selectively disrupt either SPAK or PP1 binding to NKCC1. We found that these peptides respectively reduce or enhance KCC2 membrane clustering and function, supporting the existence of a functional signaling complex that links the two transporters. Notably, peptide 5, a selective inhibitor of the NKCC1-SPAK interaction, and its optimized analog 5b, restored KCC2 membrane clustering, reduced seizure severity in vivo, and suppressed epileptiform discharges in human epileptic tissue. Together, these findings reveal a previously unrecognized mechanism of transporter crosstalk and highlight a promising therapeutic strategy to normalize chloride homeostasis and strengthen GABAergic inhibition in disorders characterized by altered KCC2 membrane expression.

## Results

### Membrane dynamics reveal the dynamic incorporation of KCC2 into NKCC1-enriched membrane clusters

We propose a novel role for neuronal NKCC1: by interacting with SPAK and PP1, NKCC1 may regulate the phosphorylation - and consequently the membrane stability - of KCC2, which itself lacks binding sites for SPAK or PP1. This implies a close functional and spatial relationship between KCC2 and NKCC1 at the neuronal membrane.

To investigate the interaction between these two transporters at the neuronal surface, we examined their spatial and dynamic relationship in cultured hippocampal neurons using single-particle tracking (SPT) in live cells. Specifically, we analyzed the lateral diffusion of KCC2 in relation to NKCC1 membrane clusters previously characterized ^16^. Cultures were co-transfected at 14 days in vitro (DIV) with KCC2-Flag and NKCC1-HA constructs, and the tagged proteins were revealed at 21 DIV by live-cell immunolabeling. KCC2 dynamics was tracked in real-time using videomicroscopy, by indirect coupling of anti-Flag antibodies to quantum dots, while NKCC1 clusters were visualized using a CY3-conjugated secondary antibody (see Methods). Analysis of individual trajectories revealed that KCC2 molecules located outside NKCC1 membrane clusters explored a significantly larger membrane area than KCC2 located within NKCC1 clusters (Fig. 1a). This broader exploration was associated with a higher instantaneous diffusion coefficient for isolated KCC2 compared to colocalized KCC2 molecules (Fig. 1b). In line with these observations, quantitative analyses showed reduced mobility (Fig. 1c) and increased confinement (Fig. 1d) of KCC2 within NKCC1 clusters compared to outside. Building on our previous findings that a subset of NKCC1 molecules is slowed and confined within endocytic pits at the neuronal surface ^16^, we compared KCC2 mobility within NKCC1 clusters and clathrin-coated pits in neurons co-transfected with clathrin-YFP (Fig. S1). KCC2 displayed significantly lower diffusion (Fig. S1a), reduced speed (Fig. S1b), and greater confinement (Fig. S1c) within NKCC1 clusters than within clathrin pits. Moreover, KCC2 molecules exhibited longer residence times in NKCC1 clusters than in clathrin pits (Fig. S1d), suggesting that KCC2 is recruited and stabilized within NKCC1 clusters, possibly through direct or indirect anchoring to the submembranous cytoskeleton via NKCC1.

**Fig. 1.**
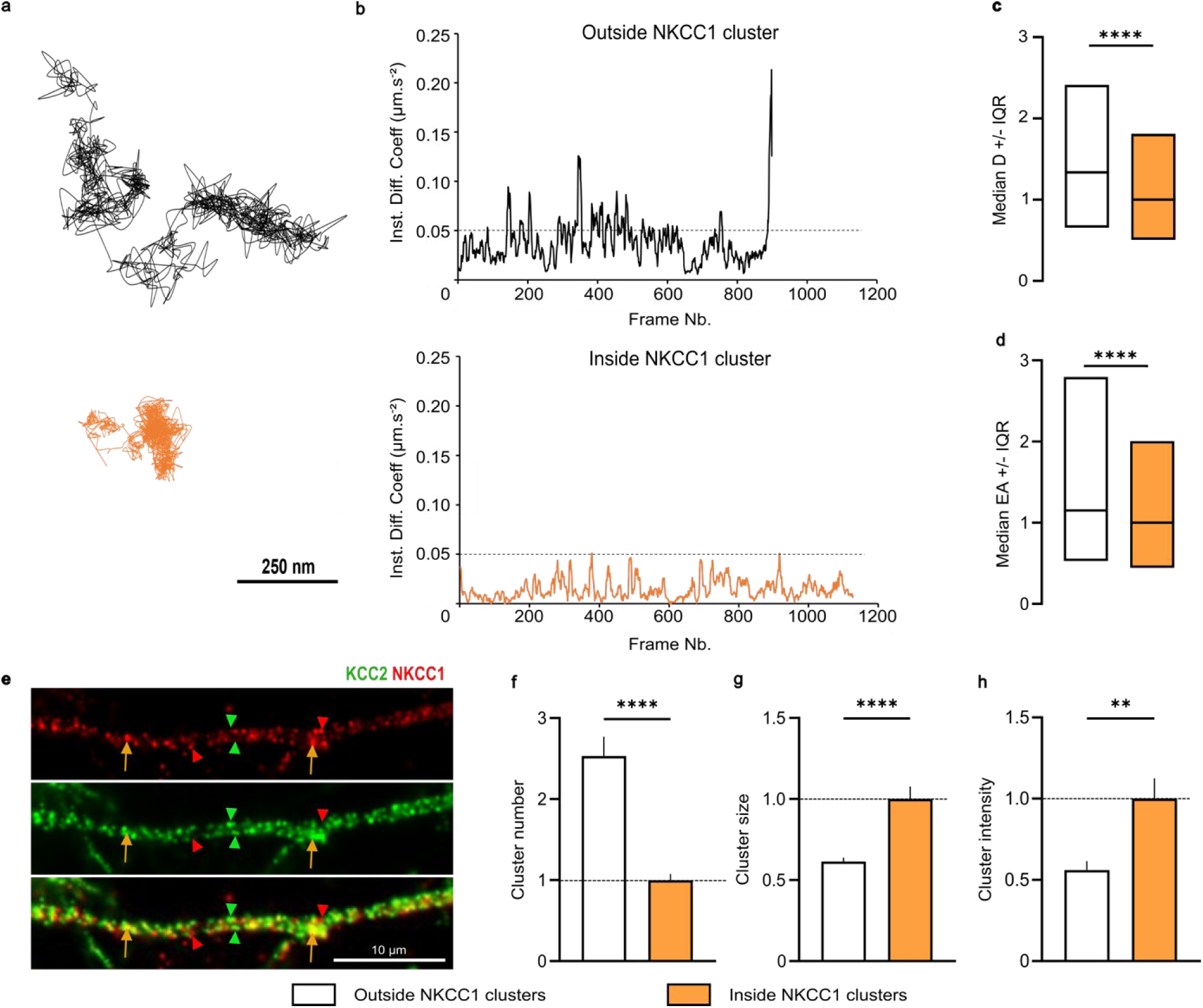
KCC2 exhibits reduced mobility and is confined within NKCC1 clusters. **a.** Examples of trajectories showing reduced surface exploration of KCC2 within NKCC1 clusters (orange) in comparison with KCC2 situated at distance from NKCC1 clusters (grey). Scale bar, 250 nm. **b.** Instantaneous diffusion coefficients corresponding to the trajectories shown in **a.** **c-d.** Median diffusion coefficient D (c) and explored area (EA) (d) ± 25–75% interquartile range (IQR) of KCC2 outside (white) or inside (orange) NKCC1 clusters. Values were normalized to the median KCC2 values inside NKCC1 clusters, 3 cultures. c, Outside: n=1148 QDs; Inside: n=343 QDs, KS test p < 0.0001; d, Outside: n=3444 QDs; Inside: n=1029 QDs, KS test p < 0.0001. **e.** Surface staining of NKCC1 (red) and KCC2 (green), in 21-DIV-old hippocampal neurons transfected with KCC2-Flag and NKCC1b-HA constructs. Orange arrows indicate KCC2 clusters that colocalize with NKCC1 clusters, while red and green arrowheads indicate NKCC1 and KCC2 clusters, respectively, without colocalization. Scale bar, 10 µm. **f-h.** Quantification of the mean number (f), size (g), and intensity (h) of KCC2 clusters that colocalize with NKCC1 clusters (orange) compared to those that do not (white). Data are shown as mean ± SEM. Values were normalized to the colocalized cluster values. n= 43 cells, 3 cultures. Cluster number: MW test, p < 0.0001; size: p < 0.0001; and intensity: p = 0.0084.

If KCC2 is indeed recruited and stabilized within NKCC1 clusters, both transporters should form joint membrane aggregates. In previous work, we showed that over 50% of NKCC1 clusters colocalize with KCC2, and that NKCC1 clusters colocalized with KCC2 are significantly more intense than isolated NKCC1 clusters, suggesting a recruitment of NKCC1 to KCC2-rich domains ^16^. Here, we explored the reverse relationship. KCC2 clusters were found to colocalize with NKCC1 clusters (Fig. 1e), although distinct clusters of either transporter could also form independently (Fig. 1e). Quantitative analysis showed that isolated KCC2 clusters were approximately 2 times more numerous than colocalized clusters (Fig. 1f). However, colocalized clusters were on average twice as large (Fig. 1g) and denser in transporter molecules (Fig. 1h) than isolated KCC2 clusters.

Altogether, our findings indicate that subsets of KCC2 and NKCC1 transporters rely on a shared molecular scaffold for their anchoring to the submembranous cytoskeleton. This co-aggregation may reflect the assembly of a signaling platform, in which NKCC1 serves to recruit SPAK to the membrane, enabling phosphorylation of KCC2, which is otherwise unable to interact with SPAK directly.

### Proof of concept: NKCC1 recruits PP1 and SPAK to regulate KCC2 membrane stability and aggregation

The phosphatase PP1 dephosphorylates KCC2, NKCC1, and SPAK ^22–24^. Since SPAK activity is phosphorylation-dependent, its dephosphorylation by PP1 inactivates it and thereby halts downstream signaling. To test the hypothesis that NKCC1 recruits PP1 or SPAK to modulate KCC2 phosphorylation - and thus its membrane stability and chloride extrusion capacity - we expressed a mutant NKCC1 lacking the _131_RFRVNF_136_ sequence (Q9QX10 (PDB)), related to the rat sequence), which corresponds to the canonical PP1-binding site (Fig. 2a). Disruption of this motif impairs the interaction between NKCC1 and PP1, resulting in increased NKCC1 phosphorylation and enhanced transporter activity ^25^. NKCC1 also contains two distinct SPAK-binding motifs within its N-terminal cytosolic region: the _76_RFx(Q)V_79_ motif and the _131_RFx(R)V_134_ motifs, corresponding to the SPAK1 and SPAK2 binding sites ^26,27^ (Fig. 2a). Notably, the consensus motifs for PP1 and SPAK2 binding partially overlap within the _131_RFRVNF_136_ sequence, highlighting a competitive interaction between PP1 and SPAK at this site ^25^. The NKCC1 mutant lacking the _131_RFRVNF_136_ sequence thus corresponds to a NKCC1-ΔSPAK2ΔPP1 mutant. Because the SPAK1 binding site remains intact in the NKCC1-ΔSPAK2ΔPP1 mutant, we hypothesized that this construct retains the ability to bind SPAK but not PP1. In this context, impaired PP1 recruitment would prevent SPAK dephosphorylation, maintaining SPAK in an active state. Consequently, SPAK could engage the SPAK1 motif and phosphorylate both NKCC1 and KCC2, thereby promoting KCC2 internalization (Fig. 2a).

**Fig. 2.**
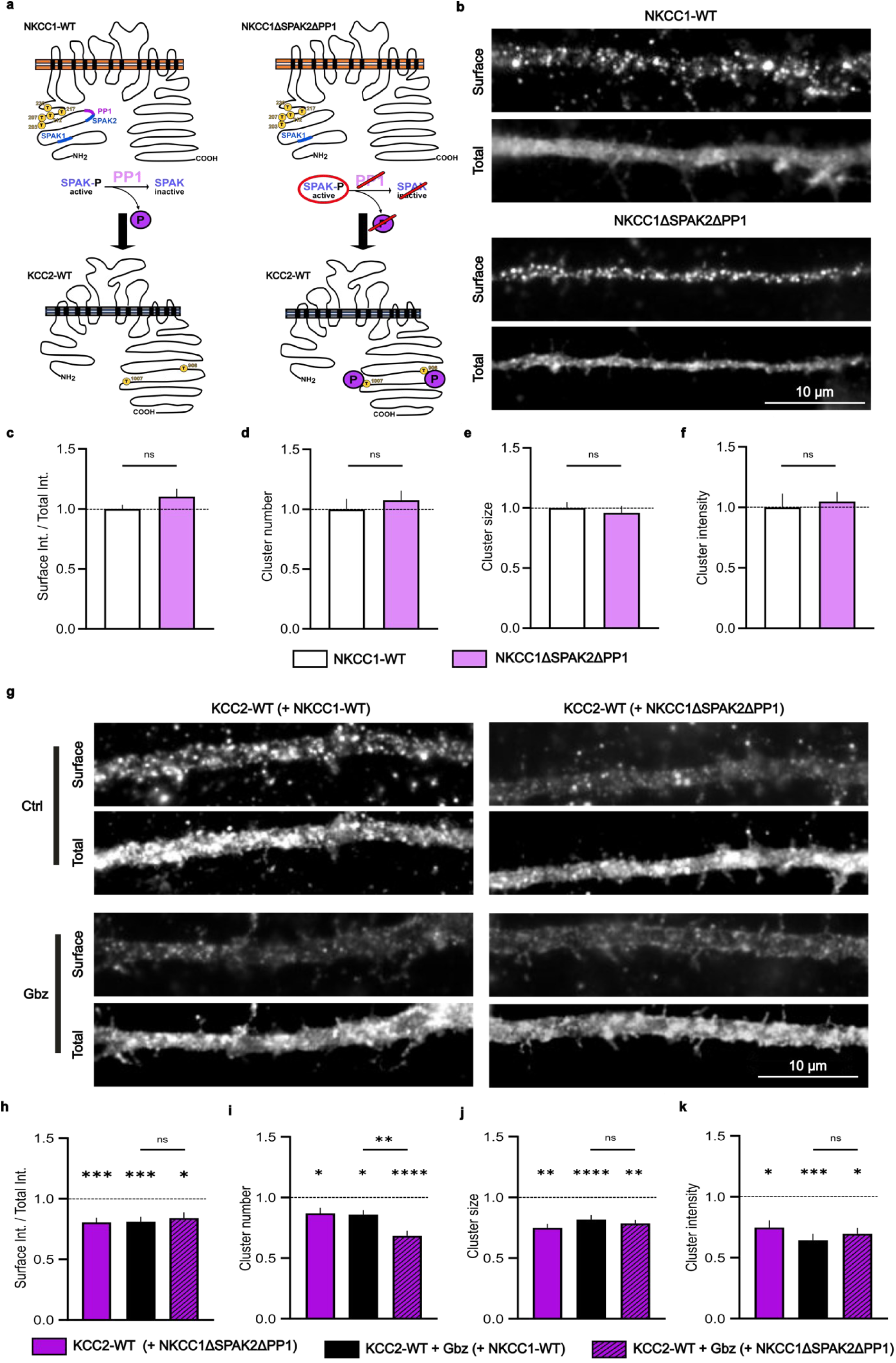
NKCC1-ΔSPAK2ΔPP1 expression reduces KCC2 membrane levels and clustering. **a.** Schematic representation of the proposed model in which NKCC1 mediates the recruitment of SPAK and PP1, regulating SPAK-dependent phosphorylation of KCC2 at threonines 906 and 1007. **b.** Surface and total (surface + intracellular) HA immunostaining of NKCC1 in neurons expressing either NKCC1-WT or NKCC1-ΔSPAK2ΔPP1. Scale bar, 10 μm. **c.** The surface-to-total pixel intensity ratio of NKCC1 remains unchanged in neurons expressing NKCC1-WT (white) or NKCC1-ΔSPAK2ΔPP1 (pink). NKCC1-WT: n = 109 cells; NKCC1-ΔPP1ΔSPAK1: n = 101 cells; 2 independent cultures. Mann–Whitney (MW) test: p = 0.9692. **d-f.** No change in the mean cluster number (d), size (e), or intensity (f) of NKCC1 clusters in neurons expressing NKCC1-WT (white) or NKCC1-ΔSPAK2ΔPP1 (pink). NKCC1-WT n = 50 cells; NKCC1-ΔSPAK2ΔPP1 n = 44 cells; 2 cultures. MW test — cluster number: p = 0.2273; cluster size: p = 0.6066; cluster intensity: p = 0.1059. **g.** Surface and total (surface + intracellular) Flag immunostaining in neurons co-expressing KCC2 and either NKCC1-WT or NKCC1-ΔSPAK2ΔPP1 under control or gabazine (Gbz) conditions. Scale bar, 10 μm. **h.** Reduced surface-to-total pixel intensity ratio of KCC2 in neurons expressing NKCC1-ΔSPAK2ΔPP1 (purple) or in neurons expressing NKCC1-WT and exposed to gabazine (black). Note that neurons expressing NKCC1-ΔSPAK2ΔPP1 do not respond to gabazine treatment (hatched purple). Values were compared to KCC2-WT + NKCC1-WT control condition. KCC2-WT + NKCC1-WT ± Gbz: Ctrl, n = 52 cells; Gbz, n = 46 cells; 3 cultures. MW test: p = 0.0005. KCC2-WT + NKCC1-WT vs KCC2-WT + NKCC1-ΔSPAK2ΔPP1: WT, n = 52 cells; ΔSPAK2ΔPP1, n = 61 cells; 3 cultures. MW test: p = 0.0003. KCC2-WT + NKCC1-WT vs KCC2-WT + NKCC1-ΔSPAK2ΔPP1 + Gbz: WT, n = 35 cells; ΔSPAK2ΔPP1 + Gbz, n = 43 cells; 2 cultures. MW test: p = 0.0238. **i-k.** Reduced mean KCC2 cluster number (d), size (e), and intensity (f) in neurons expressing expressing NKCC1-ΔSPAK2ΔPP1 (purple) or in neurons expressing NKCC1-WT and exposed to gabazine (black). Note that neurons expressing NKCC1-ΔSPAK2ΔPP1 respond less to gabazine treatment (hatched purple). Values were compared to KCC2-WT + NKCC1-WT control condition. KCC2-WT + NKCC1-WT ± Gbz: Ctrl, n = 52 cells; Gbz, n = 46 cells; 3 cultures. MW test: p = 0.0005. KCC2-WT + NKCC1-WT vs KCC2-WT + NKCC1-ΔSPAK2ΔPP1: WT, n = 52 cells; ΔSPAK2ΔPP1, n = 61 cells; 3 cultures. MW test: p = 0.0003. KCC2-WT + NKCC1-WT vs KCC2-WT + NKCC1-ΔSPAK2ΔPP1 + Gbz: WT, n = 35 cells; ΔSPAK2ΔPP1 + Gbz, n = 43 cells; 2 cultures. MW test: p = 0.0238 KCC2-WT + NKCC1-WT ± Gbz: Ctrl, n = 38 cells; Gbz, n = 40 cells; 3 cultures. MW test — cluster number: p = 0.0299; cluster size: p = 0.0077; cluster intensity: p = 0.0457. KCC2-WT + NKCC1-WT vs KCC2-WT + NKCC1-ΔSPAK2ΔPP1: WT, n = 62 cells; ΔSPAK2ΔPP1, n = 55 cells; 3 cultures. MW test — cluster number: p = 0.0475; cluster size: p < 0.0001; cluster intensity: p = 0.0004. KCC2-WT + NKCC1-WT vs KCC2-WT + NKCC1-ΔSPAK2ΔPP1 + Gbz: WT, n = 26 cells; ΔSPAK2ΔPP1 + Gbz, n = 38 cells; 2 independent cultures. MW test — cluster number: p < 0.0001; cluster size: p = 0.0014; cluster intensity: p = 0.0365. All data are presented as mean ± SEM. Values were normalized to the respective NKCC1-WT (c-f) or KCC2-WT + NKCC1-WT (h-k) control values.

While the WNK/SPAK pathway is known to stabilize NKCC1 at the membrane through phosphorylation-dependent mechanisms ^28,29^, we previously demonstrated that pharmacological inhibition of the pathway with WNK463 or closantel - selective inhibitors of WNK1 and SPAK, respectively - has little effect on NKCC1 membrane or total expression in hippocampal neurons under basal activity conditions ^17^. Only under hyperactive conditions is a significant increase in the surface-to-total NKCC1 ratio observed, which is blocked by WNK pathway inhibition ^17^. Consistent with these findings, under conditions of excitatory transmission silencing with tetrodotoxin (TTX, 1 μM), kynurenic acid (KYN, 1 mM), and the group I/II mGluR antagonist R,S-MCPG (500 μM), expression of NKCC1-ΔSPAK2ΔPP1 did not alter NKCC1-HA immunoreactivity before or after permeabilization, indicating that both surface and total NKCC1 levels remained unchanged (Fig. 2b). Quantitative analyses confirmed comparable surface/total NKCC1 ratios between NKCC1-ΔSPAK2ΔPP1 and NKCC1-WT expressing neurons (Fig. 2c). Moreover, the mutant had no effect on NKCC1 cluster number (Fig. 2d), size (Fig. 2e), or intensity (Fig. 2f), indicating that membrane aggregation of NKCC1 remains unaffected under basal conditions and is PP1 independent. In contrast, expression of NKCC1-ΔSPAK2ΔPP1 significantly reduced KCC2 immunoreactivity, particularly at the cell surface, under basal conditions (Fig. 2g). Quantification revealed a significant decrease in the surface/total KCC2 ratio in NKCC1-ΔSPAK2ΔPP1-expressing neurons compared to NKCC1-WT controls (Fig. 2h). This decrease in KCC2 membrane stability was accompanied by a significant reduction in its membrane clustering, as reflected by a decreased number of clusters (Fig. 2i), as well as reduced cluster size (Fig. 2j) and intensity (Fig. 2k).

The WNK/SPAK pathway is activated by reductions in intracellular chloride. In cultured neurons, chloride influx occurs primarily through GABA_A_ receptors. Blocking these receptors with the selective antagonist gabazine activates the WNK/SPAK pathway, leading to KCC2 phosphorylation, membrane declustering, and internalization ^15^. We compared the effect of acute gabazine treatment on KCC2 expression in neurons transfected with either NKCC1-WT or NKCC1-ΔSPAK2ΔPP1. Gabazine exposure led to a marked reduction in KCC2 surface expression (Fig. 2g-h) and clustering in NKCC1-WT-expressing neurons, evidenced by decreased cluster number (Fig. 2i), size (Fig. 2j), and intensity (Fig. 2k). In contrast, gabazine had no additive effect on surface KCC2 levels in neurons expressing NKCC1-ΔSPAK2ΔPP1 (Fig. 2h). However, it further reduced the number of KCC2 clusters (Fig. 2i), while cluster size and intensity remained unaffected (Fig. 2j-k). Thus, expressing a mutant NKCC1 incapable of recruiting PP1 and inactivating SPAK impacts KCC2 surface expression and membrane clustering. This effect resembles that observed following pharmacological activation of the WNK/SPAK pathway with gabazine ^15^. The additional reduction in KCC2 cluster number in gabazine-treated NKCC1-ΔSPAK2ΔPP1 neurons suggests that SPAK can still be further activated in this context.

Collectively, these results provide a proof of concept that NKCC1 contributes to the membrane regulation of KCC2 by recruiting both PP1 and SPAK.

### Development of SPAK-activating peptides to reduce membrane KCC2 and elevate intracellular chloride levels

To date, no pharmacological activators of SPAK have been identified. While SPAK inhibition has been a major therapeutic focus for counteracting the loss of KCC2 and the increased NKCC1 surface expression observed in many neurological disorders ^3–9^, elevated KCC2 surface expression has also been documented in human intractable temporal lobe epilepsy ^10^ and in inflammatory pain conditions such as acute arthritis ^11^. In these contexts, activating SPAK - by promoting KCC2 membrane destabilization - could therefore represent a novel therapeutic strategy.

Our findings using the NKCC1-ΔSPAK2ΔPP1 mutant further suggest that a peptide mimicking the PP1-binding motif on NKCC1 could be developed to competitively block PP1 recruitment. Such a strategy would indirectly enhance SPAK activity by preventing PP1-mediated dephosphorylation of nearby KCC2 and NKCC1 co-transporters.

Using protein-protein interaction modeling, we designed several candidate peptides (peptides 1, 2, 3 and 4) derived from distinct sequences surrounding the RFRVNF motif of NKCC1 (Fig. 3 a-b). Each peptide was fused to a TAT sequence to facilitate cell permeability (see Methods). To validate cellular uptake, we first tested a biotinylated control TAT peptide. Following 1-hour incubation at 37 °C in culture, streptavidin-based fluorescent labeling revealed robust internalization of the peptide throughout the neuronal soma and processes, confirming efficient cellular penetration (Fig. S2a).

**Fig. 3.**
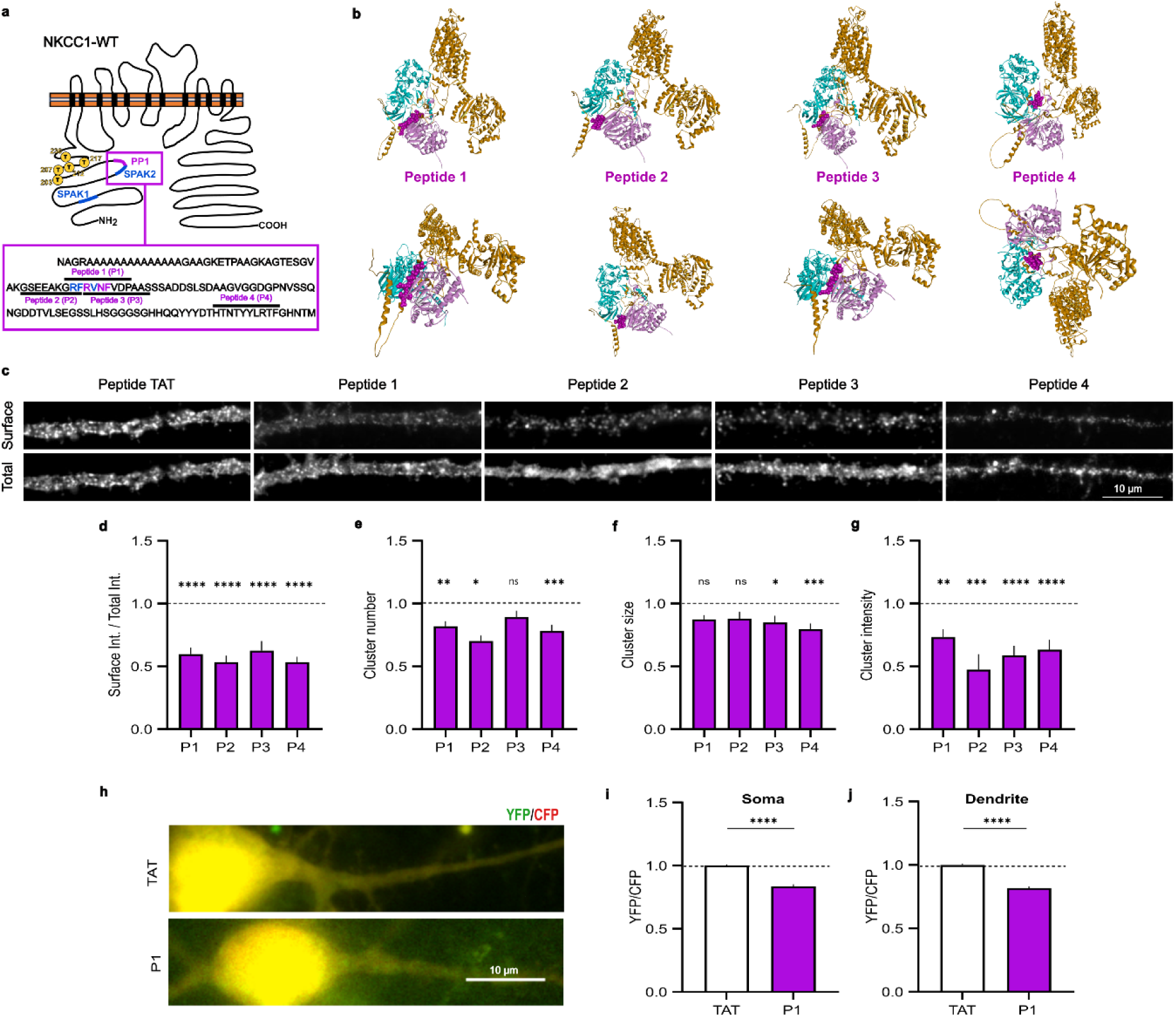
Peptides 1–4 activate SPAK and reduce KCC2 membrane clustering, with peptide 1 also increasing intracellular chloride. **a.** Schematic representation of NKCC1 showing peptides 1, 3, and 4, along with the positions they mimic. **b.** Structure of NKCC1 (brown), SPAK (blue), and PP1 (pink). The purple regions correspond to peptide sequences 1, 3 and 4. Top and bottom: different angles of the molecular complex are shown. **c.** Surface and total (surface + intracellular) Flag immunostaining of KCC2 in neurons co-expressing KCC2 and NKCC1, treated for 1 hour with control peptide (TAT) or peptide 1 (P1), peptide 3 (P3), or peptide 4 (P4) at 100 µM. Scale bar, 10 μm. **d.** Reduced surface-to-total pixel intensity ratio of KCC2 (purple) in neurons co-expressing NKCC1 following exposure to peptide 1 (P1), peptide 3 (P3), or peptide 4 (P4). P1: TAT, n = 67 cells; P1, n = 67 cells; 3 cultures; MW test, p < 0.0001. P2: TAT, n = 67 cells; P1, n = 70 cells; 3 cultures; MW test, p < 0.0001. P3: TAT, n = 39 cells; P3, n = 35 cells; 2 cultures; MW test, p < 0.0001. P4: TAT, n = 72 cells; P4, n = 72 cells; 3 cultures; MW test, p < 0.0001. **e-g.** Reduced mean KCC2 cluster number (d), size (e), and intensity (f) in neurons expressing NKCC1 following treatment with peptides P1, P3, or P4. P1: TAT, n = 73 cells; P1, n = 68 cells; 5 cultures; MW test — cluster number: p = 0.0018; cluster size: p = 0.0793; cluster intensity: p = 0.0053. P2: TAT, n = 21 cells; P2, n = 17 cells; 2 cultures; MW test — cluster number: p = 0.0124; cluster size: p = 0.2909; cluster intensity: p = 0.0003. P3: TAT, n = 34 cells; P3, n = 34 cells; 2 cultures; MW test — cluster number: p = 0.1888; cluster size: p = 0.0444; cluster intensity: p < 0.0001. P4: TAT, n = 44 cells; P4, n = 49 cells; 3 cultures; MW test — cluster number: p = 0.0005; cluster size: p = 0.0002; cluster intensity: p < 0.0001. **h.** Overlay images of CFP (red) and YFP (green) in neurons expressing SuperClomeleon along with NKCC1 and KCC2, under control (TAT) and peptide 1 (P1) conditions. **i-j.** YFP/CFP fluorescence ratios in neurons expressing SuperClomeleon, NKCC1, and KCC2 under control (white) and peptide 1 (purple) conditions, measured in the soma (h) and dendrite (i). TAT: n = 54 cells; P1: n = 40 cells; MW test, p < 0.0001. In all graphs, values were normalized to their respective TAT control values.

We then evaluated the effect of the control TAT peptide on KCC2 in hippocampal neurons co-transfected with KCC2-Flag and NKCC1-HA-WT. Compared to untreated cells, TAT peptide treatment did not significantly alter the KCC2 surface-to-total expression ratio (Fig. S2 b-c), although it slightly reduced the number of KCC2 membrane clusters (Fig. S2d) without affecting their size (Fig. S2e) or fluorescence intensity (Fig. S2f). All subsequent experiments were therefore analyzed relative to the TAT control peptide.

Compared to the TAT control, peptides 1, 2, 3 and 4 markedly decreased KCC2 surface immunoreactivity in NKCC1-WT-expressing neurons (Fig. 3c), with quantification showing ∼50% reduction in the surface-to-total KCC2 ratio (Fig. 3d). This reduction in membrane stability was accompanied by a significant decrease in KCC2 clustering, as reflected by fewer clusters (Fig. 3e), reduced cluster size (Fig. 3f), and lower fluorescence intensity (Fig. 3g). These results closely resemble those observed in neurons expressing the NKCC1-ΔSPAK2ΔPP1 mutant (Fig. 2 g-k). Notably, peptides 1, 3 and 4 had a markedly reduced effect on KCC2 membrane expression (Fig. S3 a-b) and clustering (Fig. S3 c-e) in neurons expressing NKCC1-ΔSPAK2ΔPP1, indicating that their action is mediated by preventing PP1 binding to NKCC1. If these peptides prevent PP1 binding to NKCC1, thereby sustaining SPAK activation and promoting KCC2 phosphorylation, internalization, and functional downregulation, they should also impair neuronal chloride extrusion. To test this, we used SuperClomeleon imaging to monitor intracellular chloride levels. Neurons co-expressing NKCC1, KCC2, and the SuperClomeleon probe (Fig. 3h) exhibited significantly decreased YFP/CFP ratios in both somata (Fig. 3i) and dendrites (Fig. 3j) after treatment with peptide 1 relative to the TAT control, indicating elevated intracellular chloride.

The convulsant 4-aminopyridine (4-AP) is known to increase KCC2 mobility and promote its declustering and internalization, leading to elevated intracellular chloride levels - an effect that is partly dependent on the WNK/SPAK signaling pathway ^15^. Consistent with this, we observed a significant decrease in the YFP/CFP ratio of the SuperClomeleon sensor following 4-AP treatment, indicative of increased intracellular chloride (Fig. S4 a-c). Interestingly, pretreatment with peptide 1 further enhanced the 4-AP-induced chloride accumulation (Fig. S4 a-c), supporting the idea that peptide 1 amplifies the effect of 4-AP in neurons where KCC2 clustering and chloride transport function have already been compromised.

Altogether, these findings indicate that peptides 1, 2, 3, and 4 function as SPAK activators by mimicking the PP1-binding motif of NKCC1, thereby promoting KCC2 membrane destabilization and compromising chloride extrusion.

### Development of a SPAK-inhibitory peptide to enhance KCC2 membrane stability and reduce intracellular chloride levels

Reduced KCC2 membrane stability is implicated in several epilepsies, as well as in autism spectrum disorders, schizophrenia, and neuropathies ^30^. Given the elevated WNK/SPAK pathway activity reported in these conditions ^29^, inhibiting this pathway may be beneficial. We therefore sought to develop a SPAK-inhibiting peptide.

Since the SPAK2 binding site overlaps with that of PP1, we focused on generating peptides (peptides 5 and 6) targeting the distinct SPAK1 binding domain (Fig. 4 a-b). We found that peptide 5 increased KCC2 surface immunoreactivity, whereas peptide 6 decreased it (Fig. 4c). Quantification of the surface-to-total ratio showed that peptide 5 had no significant effect on KCC2 surface expression, while peptide 6 significantly reduced it (Fig. 4d). Peptide 5, however, robustly increased the number (Fig. 4e), size (Fig. 4f), and intensity (Fig. 4g) of KCC2 clusters. In contrast, peptide 6 produced a mixed effect, increasing KCC2 cluster number and size (Fig. 4 e-f), but reducing their intensity by 50% (Fig. 4g).

**Fig. 4.**
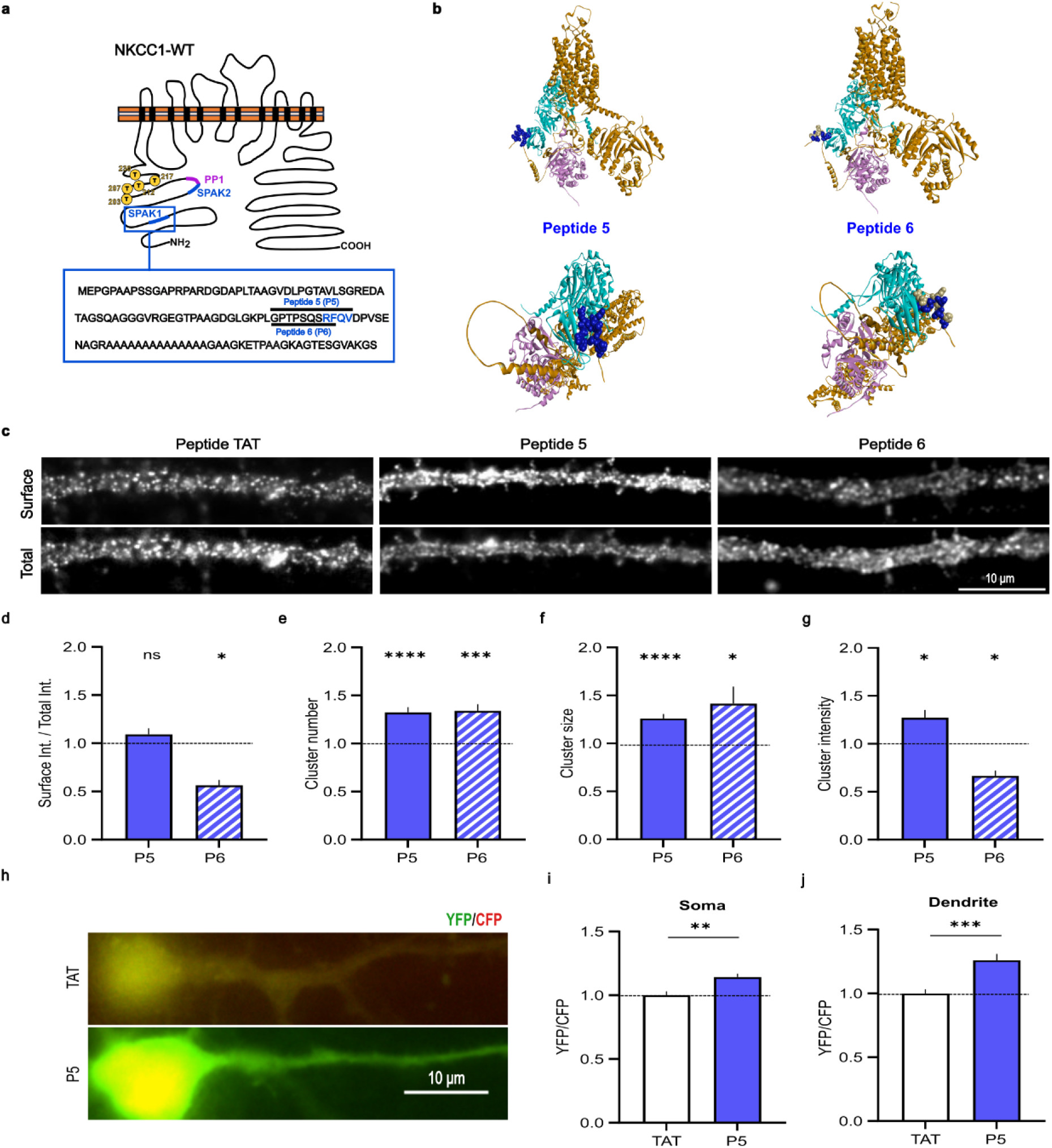
Peptide 5 inhibits SPAK, enhances KCC2 membrane clustering, and lowers intracellular chloride. **a.** Schematic representation of NKCC1 showing peptides 5 (P5) and 6 (P6), their mimicked positions, and amino acid sequences. **b.** Structure of NKCC1 (brown), SPAK (blue), and PP1 (pink). The dark blue regions correspond to peptide sequences 5 and 6. Top and bottom: different angles of the molecular complex are shown. **c.** Surface and total (surface + intracellular) Flag immunostaining of KCC2 in neurons co-expressing NKCC1 and KCC2 following 1-hour treatment with control peptide (TAT) or peptides P5 and P6 (100 μM). Scale bar, 10 μm. **d.** Surface-to-total pixel intensity ratio of KCC2 after peptide treatment. P5: TAT, n = 72 cells; P5, n = 79 cells; 5 cultures. MW test: p = 0.1834. P6: TAT, n = 40 cells; P6, n = 36 cells; 3 cultures. MW test: p = 0.0483. **e–g.** Increased mean KCC2 cluster number (d), size (e), and intensity (f) in neurons treated with peptide P5 (blue) or P6 (light blue) compared to control TAT condition (white). P5: TAT, n = 80 cells; P5, n = 90 cells; 5 cultures; MW test — cluster number: p < 0.0001; cluster size: p < 0.0001; cluster intensity: p = 0.0176. P6: TAT, n = 47 cells; P6, n = 79 cells; 3 cultures; MW test — cluster number: p = 0.0004; cluster size: p = 0.0108; cluster intensity: p = 0.0126. **h.** Overlay images of CFP (red) and YFP (green) in neurons expressing SuperClomeleon together with NKCC1 and KCC2 under control (TAT) and peptide 5 (P5) conditions. Scale bar, 10 μm. **i–j.** YFP/CFP fluorescence ratio in the soma (h) and dendrites (i) of neurons expressing SuperClomeleon and treated with peptide 5 (blue) compared to control (white). TAT, n = 35 cells; P5, n = 45 cells; 3 cultures; MW test soma: p = 0.0011; dendrite: p = 0.0001. All data are presented as mean ± SEM. Values were normalized to their respective control (TAT) condition.

SPAK and OSR1 recognize the RFxV/I motif ^31^ but also a more variable sequence KFxI/W ^32^. We therefore examined NKCC1 for sequences resembling RFxV/I and assessed their accessibility. We thus tested peptides 7 and 8 (Fig. S5 a,b), which mimic potential SPAK consensus sites - KFGWI and KFRI - located away from the known SPAK1 and SPAK2 binding motifs on NKCC1 ^32^. Neither peptide 7 nor peptide 8 significantly enhanced KCC2 clustering in treated neurons (Fig. S5 c-g).

Thus, among the peptides tested, only peptide 5 promoted KCC2 clustering in neurons. Given its clear effect on clustering, we next examined the functional consequences of peptide 5 treatment. Chloride imaging revealed that peptide 5 increased the YFP/CFP ratio of the SuperClomeleon sensor in the soma and dendrites of transfected neurons (Fig. 4h-j), indicating a reduction in intracellular chloride levels. As peptide 5 had no effect on NKCC1 clustering under basal conditions (Fig. S6 a-c), these results suggest that peptide 5 functionally modulates chloride homeostasis by promoting KCC2 clustering at the cell membrane, thereby enhancing chloride extrusion.

The specificity of peptide 5 was confirmed by the absence of effects on membrane clustering of the GABA_A_ receptor γ2 subunit - both at and outside inhibitory synapses - and on gephyrin, the key scaffolding protein for GABA_A_ receptors (Fig. S7 a-d), indicating a selective action on KCC2 without off-target effects on other critical components of inhibitory synapses. Furthermore, peptide 5 retained its ability to promote KCC2 clustering in neurons expressing the NKCC1-ΔSPAK2ΔPP1 mutant (Fig. 5 a-b), indicating that its action is not mediated by preventing PP1 binding to NKCC1, as is the case for peptides 1-4, but rather by blocking SPAK binding to NKCC1 at the SPAK1 site. Importantly, peptide 5 had no effect in neurons transfected with a phospho-mimetic KCC2 mutant at Thr906 and Thr1007, the known SPAK phosphorylation sites (Fig. 5 c-f), supporting the conclusion that the peptide acts by preventing SPAK recruitment to NKCC1 and subsequent SPAK-mediated phosphorylation of KCC2 at Thr906 and Thr1007.

**Fig. 5.**
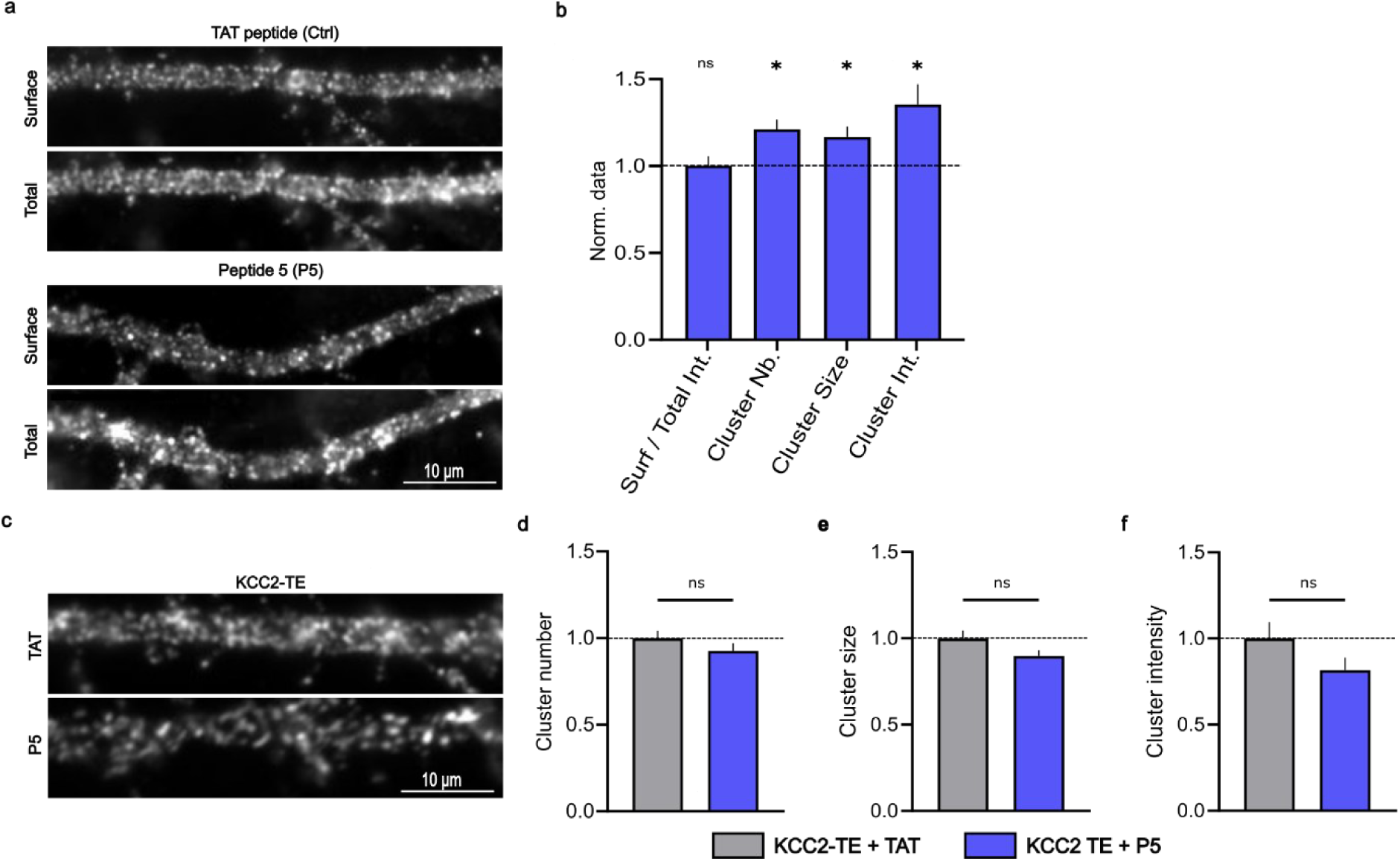
Peptide 5 inhibits SPAK binding to NKCC1, rather than PP1, and modulates KCC2 phosphorylation at Thr906 and Thr1007. **a.** Surface and total (surface + intracellular) Flag immunostaining of KCC2 in neurons co-expressing KCC2 and NKCC1-ΔSPAK2ΔPP1 following 1-hour treatment with peptides TAT or P5 (100 μM). Scale bar, 10 μm. **b.** Quantification of KCC2 surface expression and clustering in neurons co-expressing KCC2 and NKCC1-ΔSPAK2ΔPP1 following 1-hour treatment with peptides TAT or P5 (100 μM). Surface-to-total pixel intensity ratio : TAT, n = 46 cells; P5, n = 35 cells; 3 cultures; MW test: p = 0.4915. Cluster analysis: TAT, n = 38 cells; P5, n = 34 cells; 3 cultures; MW test — cluster number: p = 0.0122; cluster size: p = 0.0185; cluster intensity: p = 0.0122. **c.** Surface Flag immunostaining of the phosphomimetic mutant KCC2-T906E/T1007E (KCC2-TE) in neurons treated for 1 hour with peptides TAT or P5. Scale bar, 10 μm. **d–f.** Mean KCC2 cluster number (d), size (e), and intensity (f) in neurons expressing KCC2-TE, treated with peptides TAT or P5. KCC2-TE: TAT, n = 45 cells; P5, n = 43 cells; 3 cultures; MW test — cluster number: p = 0.4345; cluster size: p = 0.1145; cluster intensity: p = 0.1718. All data are presented as mean ± SEM. Values were normalized to their respective control (TAT) condition.

Altogether, these findings indicate that peptide 5 functions as a SPAK inhibitor by blocking its recruitment to NKCC1, thereby preventing phosphorylation of KCC2, promoting its stabilization in membrane-bound clusters, and enhancing chloride extrusion from neurons.

### Efficacy of peptide 5 under pathological conditions

Given that peptide 5 promotes KCC2 clustering and neuronal chloride homeostasis under basal activity conditions, we next investigated whether it could prevent KCC2 declustering induced by hyperexcitability and exert beneficial effects in epilepsy.

To allow for in vivo use, peptide 5 was modified into a more stable variant, peptide 5b. Due to its structural similarity to an endogenous NKCC1 sequence, peptide 5 is likely to be susceptible to degradation by peptidases. To enhance its enzymatic stability, we introduced minor amino acid substitutions by replacing threonine residues with serine and vice versa, which are structurally similar and thus unlikely to affect the overall conformation or function of the peptide (Fig. S8a). Following in vitro validation of its ability to promote KCC2 membrane clustering under basal conditions (Fig. S8 b-e), peptide 5b was tested in cultured neurons exposed to the convulsant 4-aminopyridine (4-AP).

Neurons were pretreated for 1 hour with either a control TAT peptide or peptide 5b prior to 30-minute incubation with 4-AP in the continued presence of the peptides. Cultures were then fixed and KCC2 distribution was assessed via immunostaining of the Flag epitope (Fig. 6a). When data were averaged per cell, neither 4-AP alone nor peptide 5b in the presence of 4-AP significantly affected KCC2 cluster intensity (Fig. 6 b-c). However, there was a trend toward reduced KCC2 clustering following 4-AP treatment, which appeared to be prevented by peptide 5b, as observed in both the cumulative distribution (Fig. 6b) and the mean intensity values (Fig. 6c) of cluster intensities. To determine whether this effect was specific to a subset of clusters, we performed a cluster-level analysis. 4-AP had no significant effect on the intensity of all KCC2 clusters (including those colocalized and not colocalized with NKCC1; Fig. 6 d-e) or on clusters distant from NKCC1 (Fig. 6 f-g). In contrast, 4-AP selectively reduced the intensity of KCC2 clusters colocalized with NKCC1 (Fig. 6 h-i). Notably, pretreatment with peptide 5b prevented this 4-AP-induced reduction in transporter levels within these clusters (Fig. 6 h-i).

**Fig. 6.**
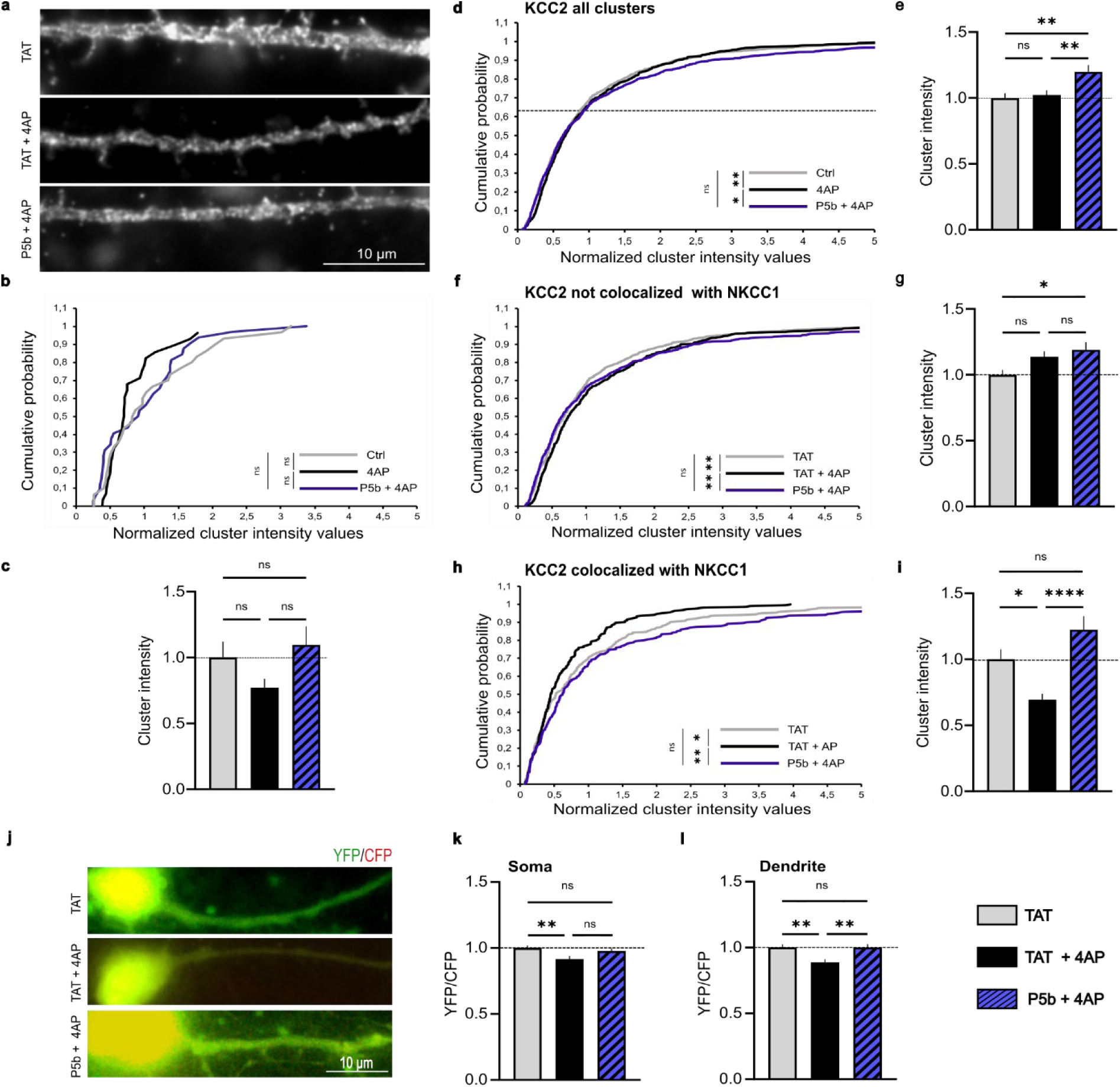
Peptide 5b prevents 4-AP–induced reduction of KCC2 clustering at NKCC1 sites and the associated elevation of intracellular chloride. **a.** Surface staining of KCC2 in 21-DIV hippocampal neurons transfected with KCC2-Flag and NKCC1b-HA constructs and treated with TAT peptides alone, TAT + 4-AP, or peptide 5b (P5b) + 4-AP. Scale bar, 10 µm. **b–c.** Cell-level analysis. Cumulative probability distributions (b) and mean intensity values (c) of KCC2 clusters (colocalized and not colocalized with NKCC1) following treatment with control peptide (TAT, grey), TAT + 4-AP (black), or P5b + 4-AP (blue). **b.** TAT: n = 32 cells; TAT + 4-AP: n = 28 cells; P5b + 4-AP: n = 30 cells; 3 cultures. Kolmogorov–Smirnov (KS) test: TAT vs TAT + 4-AP, p = 0.0838; TAT + 4-AP vs P5b + 4-AP, p = 0.1247; TAT vs P5b + 4-AP, p = 0.4866. **c.** One-way ANOVA: TAT vs TAT + 4-AP, p = 0.4260; TAT + 4-AP vs P5b + 4-AP, p = 0.1573; TAT vs P5b + 4-AP, p = 0.9110. **d–i.** Cluster-level analysis of KCC2 clusters: all clusters (d–e), clusters not colocalized with NKCC1 (f–g), and clusters colocalized with NKCC1 (h–i). Cumulative probability distributions (d, f and h) and mean intensity values (e, g and i) are shown for each condition. All clusters: **d.** TAT: n = 998; TAT + 4-AP: n = 876; P5b + 4-AP: n = 947; 3 cultures. KS test: TAT vs TAT + 4-AP, p = 0.0058; TAT + 4-AP vs P5b + 4-AP, p = 0.0158; TAT vs P5b + 4-AP, p = 0.1871. **e.** One-way ANOVA: TAT vs TAT + 4-AP, p = 0.9694; TAT + 4-AP vs P5b + 4-AP, p = 0.0091; TAT vs P5b + 4-AP, p = 0.0015. Not colocalized clusters: **f.** TAT: n = 691; TAT + 4-AP: n = 652; P5b + 4-AP: n = 706; 3 cultures. KS test: TAT vs TAT + 4-AP, p = 0.0040; TAT + 4-AP vs P5b + 4-AP, p = 0.0029; TAT vs P5b + 4-AP, p = 0.3073. **g.** One-way ANOVA: TAT vs TAT + 4-AP, p = 0.1293; TAT + 4-AP vs P5b + 4-AP, p = 0.8211; TAT vs P5b + 4-AP, p = 0.0138. Colocalized clusters: **h.** TAT: n = 307; TAT + 4-AP: n = 224; P5b + 4-AP: n = 241; 3 cultures. KS test: TAT vs TAT + 4-AP, p = 0.029; TAT + 4-AP vs P5b + 4-AP, p = 0.0064; TAT vs P5b + 4-AP, p = 0.1693. **i.** One-way ANOVA: TAT vs TAT + 4-AP, p = 0.0188; TAT + 4-AP vs P5b + 4-AP, p < 0.0001; TAT vs P5b + 4-AP, p = 0.1153. **j–l.** Peptide 5b prevents 4-AP–induced elevation of intracellular chloride in neurons. **j.** Representative overlay images of CFP (red) and YFP (green) in neurons expressing SuperClomeleon, NKCC1, and KCC2 under TAT (control), TAT + 4-AP, and P5b + 4-AP conditions. Scale bar, 10 μm. **k–l.** YFP/CFP fluorescence ratio measured in the soma (**k**) and dendrite (**l**) of neurons expressing SuperClomeleon, NKCC1, and KCC2 under TAT (grey), TAT + 4-AP (black), and P5b + 4-AP (hatched blue) conditions. TAT: n = 38 cells; TAT + 4-AP: n = 38 cells; P5b + 4-AP: n = 28 cells; 3 cultures. One-way ANOVA: Soma: TAT vs TAT + 4-AP, p = 0.0023; TAT + 4-AP vs P5b + 4-AP, p = 0.1282; TAT vs P5b + 4-AP, p = 0.8282. Dendrite : TAT vs TAT + 4-AP, p = 0.0074; TAT + 4-AP vs P5b + 4-AP, p = 0.0047; TAT vs P5b + 4-AP, p > 0.9999. All values were normalized to the corresponding TAT control condition.

These findings report that 4-AP selectively disrupts membrane clustering of KCC2 in regions where it is colocalized with NKCC1, and that the proximity of the two transporters may play a role in both the regulation of KCC2 and the protective effect of peptide 5b. Importantly, this protective effect of peptide 5b translated into a functional consequence: it blocked the 4-AP-induced increase in intracellular chloride levels (Fig. 6 j-l). Thus, peptide 5b is effective in vitro in preventing both the membrane disorganization of KCC2 and its associated loss of function under conditions of hyperactivity.

Encouraged by these results, we next tested the efficacy of peptide 5b in vivo in a pentylenetetrazole (PTZ) model of acute epileptic seizure in adult mice, which is known to trigger WNK pathway activation and hyperphosphorylation of both KCC2 and NKCC1 ^15^. Using the Racine scale (Fig. 7a), we found that peptide 5b significantly reduced seizure severity between 15 and 55 minutes after PTZ injection (Fig. 7 b-d). The proportion of animals reaching stage 5 (status epilepticus, SE) was lower in the peptide 5b-treated group compared to saline controls (Fig. 7e), and the duration of SE was significantly shortened following peptide 5b injection (Fig. 7f). These results indicate that peptide 5b decreases both the frequency and severity of seizures. This anti-epileptic effect was associated with improved survival: none of the 10 animals pretreated with peptide 5b died following PTZ injection (Fig. 7g), whereas mortality was observed in saline-treated controls (Fig. 7g). We conclude that peptide 5b exerts protective effects in acute epilepsy in the PTZ mouse model.

**Figure 7.**
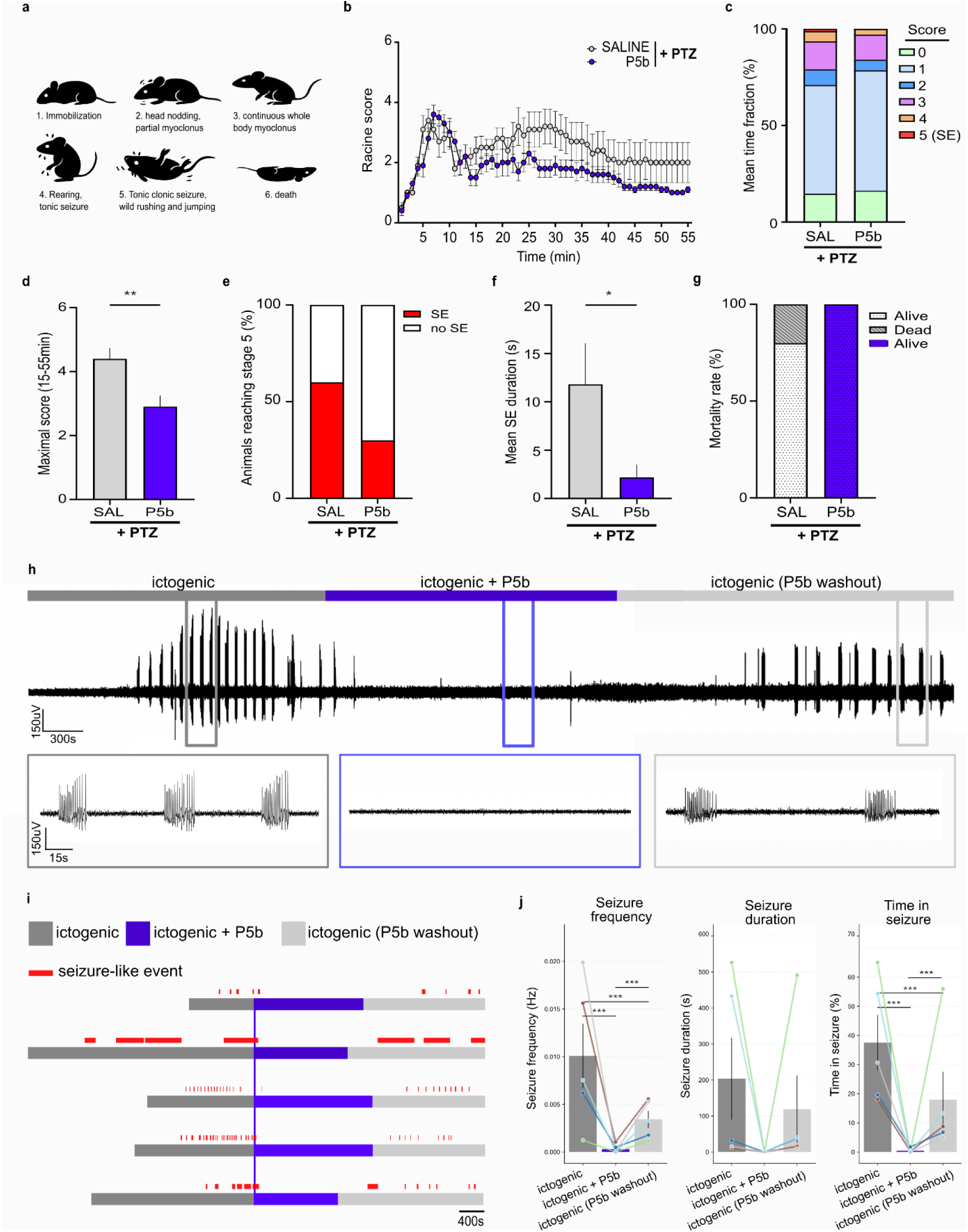
Efficacy of peptide 5b in epilepsy. **a-i.** Peptide 5b reduces acute epileptic seizure severity in the PTZ mouse model. **a.** Schematic representation of epileptic seizure severity based on Racine’s scale. **b.** Average Racine score per minute for each group: saline (grey) and peptide 5b pretreated (blue). Saline, n = 10 mice; P5b, n = 10 mice. **c.** Mean time fraction spent at each Racine stage per group (in %). **d.** Average maximal score reached per animal during the last 45 minutes. Unpaired t-test p = 0.0064 **e.** Percentage of animals reaching stage 5 in each group. **f.** Mean status epilepticus (SE) duration per group. MW test p = 0.0388. **g.** Mortality rate (in %) for each group. **h.** Voltage traces from human glioma brain slices under treatment with ictogenic conditions, after P5b treatment, and following washout. The lower panel shows enlarged views from the upper voltage trace (squares). Grey bars indicate ictogenic baseline, violet bars represent ictogenic + P5b treatment, and light grey bars represent ictogenic, P5b, washout. **i.** Timeline of the recordings for each individual human glioma brain slice color-coded according to each condition. Red lines show the seizure-like events. The vertical blue line indicates the beginning of exposure to P5b. **j.** Quantification of epileptiform activity under ictogenic conditions, after P5b treatment, and following washout. Bar plots show mean ± SEM of seizure frequency (Hz), seizure duration (s), and percentage of time in seizure across slices (n = 5). Colored dots and connecting lines indicate individual slices, highlighting paired measurements across conditions. Statistical analysis was performed using Friedman’s test for repeated measures, followed by Conover post-hoc comparisons with Holm correction. Significance levels are indicated above bars (p < 0.05, *p < 0.01, **p < 0.001).

Furthermore, we investigated the effect of peptide 5b on neuronal activity in pharmacoresistant human cortical tissue resected from patients with glioma (n = 5) using multi-electrode array (MEA) recordings. Brain slices were first exposed to an ictogenic solution to induce seizure-like activity. After seizure-like events occurred, peptide 5b was applied (ictogenic + P5b), followed by a washout with the ictogenic solution (ictogenic, P5b, washout). Strikingly, peptide 5b treatment drastically abolished seizure-like activity (Fig. 7h), an effect observed in all human brain slices tested (Fig. 7i). Quantitative analysis revealed that both seizure frequency and total time in seizure were significantly reduced in the presence of peptide 5b compared with ictogenic and washout conditions. Although seizure duration was also reduced in slices where seizures were recorded, this change did not reach statistical significance (Fig. 7j). Overall, these results indicate that peptide 5b is effective in suppressing seizure-like activity under chronic epileptic conditions in human cortical tissue.

## Discussion

We have identified a novel regulatory mechanism of the WNK/SPAK signaling pathway. Using computational modeling of protein-protein interactions, we designed several peptides to disrupt this mechanism. Peptides 1-4 act as SPAK activators, reducing KCC2 membrane clustering and chloride extrusion, whereas peptide 5 functions as a SPAK inhibitor, promoting KCC2 clustering and enhancing chloride extrusion. Notably, peptide 5 demonstrated beneficial effects in the in vivo PTZ mouse model of acute epilepsy and in pharmacoresistant human epileptic cortical tissue, underscoring its potential as a therapeutic approach for restoring inhibitory balance.

### New role of NKCC1 in mature neurons

NKCC1 expression peaks during early neuronal development and markedly declines by the end of the second postnatal week in rodents, after which it remains at consistently low levels throughout adulthood ^33^. This developmental downregulation of NKCC1 is paralleled by a robust upregulation of the K⁺-Cl⁻ cotransporter KCC2, which reaches maximal expression around postnatal day 14 and remains stable in the mature brain ^34^. As a result, mature neurons are characterized by low NKCC1 and high KCC2 expression, establishing a chloride extrusion-dominated state. This molecular switch underlies the developmental shift in GABAergic signaling: in immature neurons, high intracellular Cl⁻ maintained by NKCC1 causes GABA_A_ receptor activation to be depolarizing, whereas in mature neurons, KCC2-mediated Cl⁻ extrusion ensures that GABA evokes hyperpolarizing Cl⁻ influx ^35^. Consistent with this, multiple studies using chloride imaging and electrophysiology in neuronal cultures, brain slices, and in vivo models have shown that NKCC1 plays only a minor role in chloride homeostasis in mature neurons, where KCC2 is the dominant transporter ^36–38^.

NKCC1 is thought to play a more prominent role in non-neuronal cells than in neurons in the mature central nervous system. NKCC1 in astrocytes plays a crucial role in regulating ion homeostasis, cell volume, and buffering extracellular K⁺ and Cl⁻. It allows astrocytes to take up excess ions during neuronal activity, helping to maintain inhibitory neurotransmission and prevent neuronal hyperexcitability ^2^. In pathological conditions such as cerebral edema, trauma, and epilepsy, NKCC1 expression is upregulated in astrocytes, promoting cell swelling through osmotic water entry ^39^. Interestingly, while neuronal NKCC1 has been associated with increased seizure susceptibility, astrocytic NKCC1 appears to play a neuroprotective role by limiting network excitability ^40,41^. NKCC1 also contributes to synaptic plasticity. it has been shown to regulate astrocyte process withdrawal from synapses during long-term potentiation, thereby influencing glutamatergic transmission and modulating structural and functional plasticity at excitatory synapses ^2^. NKCC1 also regulates microglial morphology, process recruitment after injury, and ion homeostasis, and its deficiency primes microglia toward exaggerated inflammatory responses, worsening brain injury outcomes ^42^. Further, microglial NKCC1 controls interneurons neurogenesis after traumatisms ^43^.

Our findings suggest that, in mature neurons, NKCC1 assumes an unexpected, non-canonical role that goes beyond chloride transport. Rather than acting solely as an ion transporter, NKCC1 may provide a structural platform that spatially organizes SPAK and PP1, thereby orchestrating opposing phosphorylation-dephosphorylation cycles on both KCC2 and NKCC1, as well as regulating SPAK activity through PP1 ^19,44^. Such a scaffold-based mechanism positions NKCC1 as a central signaling hub capable of dynamically tuning KCC2 stability and clustering at the neuronal membrane. This perspective broadens our understanding of how inhibitory strength is regulated, and raises the possibility that targeting the NKCC1-SPAK-PP1 axis may represent a novel strategy to restore KCC2 function. The contrasting effects of peptides 1-4 and 5 on KCC2 further reinforce this model, providing proof-of-concept that selective interference with NKCC1-dependent signaling can shift the balance between destabilization and stabilization of inhibitory transmission.

An open question is whether NKCC1 is the only protein capable of recruiting both PP1 and SPAK to the membrane and of regulating the membrane aggregation and activity of KCC2. Notably, KCC2a, unlike the KCC2b isoform, possesses a SPAK-binding site in its N-terminal domain ^45^. This KCC2a isoform is predominant during early developmental stages ^2,45^. Consequently, KCC2a regulation via SPAK may not require NKCC1 scaffolding. However, KCC2a cannot bind PP1 ^19^, and thus the fine-tuning of SPAK activity via PP1-mediated dephosphorylation cannot be orchestrated by KCC2a itself alone.

### Peptides 1-4 and Peptide 5 as therapeutic tools in epilepsy and other disorders with impaired inhibition

Pharmacoresistant epilepsies, affecting around 30% of patients, represent a critical medical need for new treatments ^46,47^. These forms include development epileptic encephalopathies as well as focal onset seizures, which are often refractory to conventional therapies and frequently require alternative approaches, including surgery ^48^. Current medications affect neuronal excitability, mostly acting on ion channels, and both inhibitory and excitatory neurotransmission at postsynaptic and neurotransmitter release levels. Gene therapies are being developed to target specific mutations or modulate epileptogenic overexpression of glutamate receptors such as the kainate receptors subunit GluK2 ^49^, paving the way for more personalized medicine^50^. However, none of these treatments target KCC2 ^51^. In drug-resistant epilepsy, KCC2 function is impaired, leading to intracellular Cl⁻ accumulation and an impairment of GABA’s action ^51,52^. Some KCC2 activators and expression modulators have been developed, such as OV350, CLP257, and WNK463, showing promising results in animal models ^53–55^. However, OV350 does not affect KCC2’s membrane localization, and WNK463 presents numerous side effects and does not cross the blood-brain barrier (BBB) ^53,56^. In contrast, peptide 5 alters KCC2 membrane localization and crosses the BBB, as evidenced by its in vivo effects in mice, which distinguishes it and may enable synergistic use with the KCC2 activator OV350 for optimal modulation of KCC2 ^54^.

Furthermore, to our knowledge, no pharmacological activators of SPAK have been developed or marketed to date. Peptides 1-4 fulfill this role and thus represent novel therapeutic tools for the treatment of disorders associated with increased KCC2 membrane expression or decreased NKCC1 activity. Notably, elevated KCC2 levels have been reported in certain drug-resistant human temporal lobe epilepsy ^10^, as well as in inflammatory conditions such as acute arthritis ^11^.

Therefore, our Peptides 1-4 and Peptide 5 hold significant therapeutic potential, which has recently been protected by two patents, and future studies will determine their efficacy in treating relevant disorders.

## Online Methods

### Animals

For all experiments performed on animals, procedures were carried out according to the European Community Council directive of 24 November 1986 (86/609/EEC), the guidelines of the French Ministry of Agriculture and the Direction Departmental de la Protection des Populations de Paris (Institut du Fer à Moulin, Animalerie des Rongeurs, license C 72-05-22). All efforts were made to minimize animal suffering and to reduce the number of animals used. Timed pregnant Sprague-Dawley rats were supplied by Janvier Lab (Le Genest St. Isle, France) and embryos were used at embryonic day 18 or 19 as described below. C57BL/6JRj mice, supplied by Janvier Lab, were delivered to our animal facility at least a week before experiments. Animals were housed in standard laboratory cages on a 12-hours light/dark cycle, in a temperature-controlled room (21 °C) with free access to food and water.

### Dissociated hippocampal cultures

Primary cultures of hippocampal neurons were prepared as previously described ^13^. Briefly, hippocampi were dissected from embryonic day 18 or 19 Sprague-Dawley rats of either sex. Tissue was then trypsinized (0.25% v/v), and mechanically dissociated in 1X HBSS (Invitrogen, Cergy Pontoise, France) containing 10 mM HEPES (Invitrogen,). Neurons were plated at a density of 180 × 10^3^ cells/mL onto 18-mm diameter glass coverslips (Assistent, Winigor, Germany) pre-coated with 50 μg/mL poly-D,L-ornithine (Sigma-Aldrich, St. Louis, USA) in plating medium composed of Minimum Essential Medium (MEM, Invitrogen) supplemented with horse serum (10% v/v, Invitrogen), L-glutamine (2 mM) and Na^+^ pyruvate (1 mM) (Invitrogen). After attachment for 3–4 h, cells were incubated in culture medium that consists of Neurobasal medium (Invitrogen) supplemented with B27 (1X) (Invitrogen), L-glutamine (2 mM) (Invitrogen), and antibiotics (penicillin 200 units/mL, streptomycin, 200 μg/mL) (Invitrogen) for up to 3 weeks at 37°C in a 5% CO_2_ humidified incubator. Each week, one-third of the culture medium volume was renewed.

#### DNA constructs

The pcDNA3.1 Flag-YFP-hNKCC1b-HA-ECL2 (NT931) was a gift from Biff Forbush (Addgene plasmid # 49063; http://n2t.net/addgene:49063; RRID: Addgene_49063) ^57^. From this NKCC1-HA-Flag-mVenus plasmid, the NKCC1-HA-Δflag-ΔmVenus construct was raised by truncation of the tags located on NKCC1 N-terminal domain ^17^. This construct called NKCC1-HA-WT was used to raise the NKCC1-ΔSPAK2ΔPP1 construct which consisted in deleting the RFRVNF sequence. The following constructs were also used: pCAG_rat KCC2-3Flag-ECL2, pCAG_KCC2-3Flag-ECL2 T906/1007E ^15^, eGFP (Clontech), clathrin-YFP (kindly provided by D. Choquet, iiNS, Bordeaux, France), and SuperClomeleon ^58^ (kindly provided by G.J. Augustine, NTU, Singapore). All constructs were sequenced by Beckman Coulter Genomics (Hope End, Takeley, U.K).

#### Neuronal transfection

Neuronal transfections were carried out at DIV 13–14 using Transfectin (BioRad, Hercules, USA), according to the manufacturers’ instructions (DNA:transfectin ratio 1 µg:3 µl), with 1–2 µg of plasmid DNA per 20 mm well. The following ratios of plasmid DNA were used in co-transfection experiments: 0.4:0.4:0.4 µg for NKCC1, KCC2 constructs with eGFP; 0.7:0.7:0.7 µg for clathrin-YFP + KCC2-WT + NKCC1-WT; 0.3:0.3:0.3 µg for eGFP + KCC2-TE + NKCC1-WT ; 0.3:0.3:0.3 µg for eGFP + KCC2-WT + NKCC1-ΔSPAK2ΔPP1; 1:0.7:0.7 µg for SuperClomeleon + KCC2-WT + NKCC1-WT.

### Molecular Modelling

#### Model building

Structures of the full NKCC1 transporter was generated using AlphaFold3 (AF3) ^59^, Electron Microscopy data of the transporter in complex with bumetanide (PDB: 9C0H) ^60^, and electron microscopy data of the transporter without ligand (PDB: 7MXO) [https://doi.org/10.1126/sciadv.abq0952]. The full 3D model predicted by AF3 displayed very low to low confidence in the regions corresponding to NTD of NKCC1, To address this, we generated a 3D homology model and refined the model using classical homology modeling ^61,62^. The resulting protein structures were solvated using Biovia Discovery Studio (Dassault Systèmes), with a 9 Å water box and 0.1 M NaCl. Energy minimization was then performed using the Adopted Basis Newton-Raphson (ABNR) algorithm ^63,64^ and the Generalized Born with Implicit Membrane (GBIM) solvent model, until the root-mean-square gradient (RMSD) converged to 0.001.

#### Complex generation

SPAK and PP1 were docked independently onto its corresponding NKCC1 domain using the ZDock algorithm within Discovery Studio ^65^, which employs rigid-body docking to predict protein–protein complex formations. For each docking trial, approximately 2,000 poses were generated, then clustered based on spatial proximity and interaction energy ^66^.

To avoid steric bias introduced by pre-bound complexes, each docking simulation was conducted in independent manner, with no other interactors present. Representative poses from each cluster were typed using the CHARMm Polar H force field and subjected to further refinement, including: (i) solvation in a 9 Å water box with 0.1 M NaCl ionic strength, (ii) energy minimization using the Adopted Basis Newton-Raphson (ABNR) algorithm until convergence to a gradient threshold of 0.001, and (iii) 1 ns molecular dynamics equilibration using NAMD, all performed within the Discovery Studio environment.

The final pose from each cluster was selected based on structural stability and the quality of the protein–protein interface and peptide 1-8 were designed according to binding interface.

#### Peptides

The following peptides were designed using the BIOVIA Discovery Studio by Dassault system software: Peptide TAT control: GRKKRRQRRRPP; Peptide 1: GRKKRRQRRRPP-GRFRVNFVDP; Peptide 2: GRKKRRQRRRPP-GSEEAKGRF; Peptide 3: GRKKRRQRRRPP-RVNFVDPAAS; Peptide 4: GRKKRRQRRRPP-HTNTYYLRTF; Peptide 5: GRKKRRQRRRPP-GPTPSQSRFQV; Peptide 5b: GRKKRRQRRRPP-GPSPTQTRFQV; Peptide 6: GRKKRRQRRRPP-GPTPSQSRF; Peptide 7: GRKKRRQRRRPP-KGVVKFGWIK; Peptide 8: GRKKRRQRRRPP-LSKFRIDFSD. They were produced by the Sorbonne University platform. In addition, biotinylated TAT-peptide was kindly provided by Fabienne Burlina (LBM, Sorbonne Université, France).

### Peptide treatment and pharmacology

The following peptides and drugs were used: TAT, P1-P8 (100 µM), TTX (1 µM; Latoxan, Valence, France), R,S-MCPG (500 µM; Abcam, Cambridge, UK), S-MCPG (250 µM; HelloBio), Kynurenic acid (1 mM; Abcam), gabazine (10 µM; Abcam) and 4-Aminopyridin (100 µM; Thermo Fisher Scientific, ≠ J61470.14). R,S-MCPG and S-MCPG were prepared in equimolar concentrations of NaOH; TTX in 2% citric acid (v/v); closantel in DMSO (Sigma). Equimolar DMSO concentrations were used for control experiments in these conditions.

#### For SPT experiments

neurons were pre-incubated for one hour at 37 °C in a CO₂ incubator with the appropriate peptide (control TAT or P1–P8) added to the culture medium. They were then labeled for KCC2-Flag using quantum dots (see labeling procedure below) in the absence of peptide, transferred to a recording chamber, and incubated for 10 minutes at 31 °C in imaging medium (see composition below), this time in the presence of the peptide or the peptide + 4-aminopyridin when required. Imaging was performed within 50 minutes in the presence of the appropriate drug.

#### For immunofluorescence experiments

drugs and peptides were added directly to the culture medium for 1 hour in a CO₂ incubator at 37 °C, prior to fixation and immunocytochemistry.

#### For chloride imaging

neurons were incubated with the peptide for one hour at 37 °C in culture medium. They were then transferred to a recording chamber and incubated for 10 minutes at 31 °C in imaging medium (see composition below), still in the presence of the peptide or the peptide + 4-aminopyridin when required. Imaging was performed within 50 minutes in the presence of the appropriate drug.

#### For PTZ experiments

mice received intraveinous injection of vehicule (saline) or P5 (20mg/kg) 1-hour prior pentylenetetrazole subcutaneous injection.

#### The imaging medium

consisted of phenol red-free minimal essential medium supplemented with glucose (33 mM; Sigma) and HEPES (20 mM), glutamine (2 mM), Na^+^-pyruvate (1 mM), and B27 (1X) from Invitrogen.

### KCC2-Quantum Dot labeling for Single Particle Tracking of KCC2-Flag in relation to NKCC1 or clathrin clusters

Neurons expressing clathrin-YFP were stained with quantum dot–coupled KCC2-Flag following the protocol described previously ^15^, with some modifications to enable labeling of NKCC1 clusters using CY3-coupled antibodies, in order to track the mobility of KCC2 in relation to clathrin or NKCC1 aggregates. Briefly, cells were incubated for 6 min at 37 °C with primary antibodies against Flag (mouse, 1:300, Sigma, cat #F3165) and against HA antibody (rabbit, 1:300, Cell signaling Technology, cat #C29F4), washed, and incubated for 6 min at 37°C with biotinylated Fab secondary antibodies (goat anti-mouse: 1:700; Jackson Immunoresearch, cat #115-067-003, West Grove, USA) and CY3-conjugated donkey anti-rabbit secondary antibodies (1:300, 711-165-152, Jackson Immunoresearch) in imaging medium. After washes, cells were incubated for 1 min with streptavidin-coated quantum dots (QDs) emitting at 655nm (1nM; Invitrogen ref Q10123MP) in PBS (1M; Invitrogen) supplemented with 10% Casein (v/v) (Sigma) to prevent non-specific binding. Washing and incubation steps were done in imaging medium.

#### Single particle tracking and analysis

Cells were imaged using an Olympus IX71 inverted microscope equipped with a 60X objective (NA 1.42; Olympus, Tokyo, Japan) and an X-Cite 120Q (Excelitas Technologies Corp., Waltham, USA). Individual images of clathrin-YFP and NKCC1-HA-WT, and QD real time recordings (integration time of 30 ms over 1200 consecutive frames) were acquired with Hamamatsu ImagEM EMCCD camera (Hamamatsu, Japan) and MetaView software (Meta Imaging 7.7, Molecular Devices, USA). Cells were imaged within 50 min following labeling. QD tracking and trajectory reconstruction were performed with homemade software (MATLAB; The Mathworks, Natick, USA) as described in ^67^. One sub-regions of dendrites were quantified per cell. In cases of QD crossing, the trajectories were discarded from analysis. Trajectories were considered inside NKCC1 clusters or endocytic zones when overlapping with the mask of NKCC1-HA or clathrin-YFP or their dilated zone of 2 pixels (380 nm). Out of this vicinity, trajectories were considered outside NKCC1 clusters or endocytic zones. Values of the mean square displacement (MSD) plot versus time were calculated for each trajectory by applying the relation:

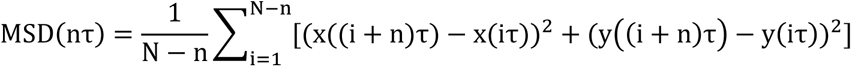

where τ is the acquisition time, N the total number of frames, n and i positive integers with n determining the time increment ^68^. Diffusion coefficients (D) were obtained by fitting the first four points without origin of the MSD vs. time curves with: MSD(nτ) = 4Dnτ + σ ; with σ being the spot localization accuracy. The explored area of each trajectory was defined as the MSD value of the trajectory at two different time intervals of at 0.42 and 0.45 s ^69^. Synaptic dwell time was defined as the duration of detection of QDs in endocytic zones or inside NKCC1 clusters on a recording divided by the number of exits as detailed previously. Dwell times ≤5 frames were not retained.

Experimenters were blind to the condition of the sample analyzed. Sample size selection for experiments was based on published experiments, pilot studies as well as in-house expertise.

#### Immunocytochemistry

After being exposed for 1 hour at 37°C to the appropriate drugs, cells were fixed for 4 min at room temperature (RT) in paraformaldehyde (PFA, 4% w/v, Sigma) and sucrose (14% w/v, Sigma) solution prepared in PBS (1X). We then stained KCC2-Flag together or not with NKCC1-HA or GABA_A_R γ2 and gephyrin following different protocols, detailed below.

Sets of neurons compared for quantification were labeled simultaneously.

##### NKCC1 surface staining

NKCC1-HA membrane expression and clustering was assessed with staining performed after a short fixation at room temperature (RT) in paraformaldehyde (PFA; 4% w/v; Sigma) and sucrose (20% w/v; Sigma) solution in 1× PBS. The cells were then washed in PBS and incubated for 30 min at RT in goat serum (GS; 10% v/v; Invitrogen) in PBS to block non-specific staining. Neurons were then incubated for 60-120 min at RT with HA antibody (rabbit, 1:500, Cell signaling Technology, cat #C29F4) in PBS–GS blocking solution. After washing, neurons were incubated with Cy™3 AffiniPure Goat Anti-Rabbit IgG (H+L) (1.5 µg/ml; Jackson ImmunoResearch, cat #111-165-003) for standard epifluorescence assays in PBS-GS solution. The coverslips were then washed, and mounted on slides with mowiol 844 (48 mg/ml; Sigma).

##### KCC2 surface staining

KCC2-Flag membrane expression and clustering was assessed with staining performed after a short fixation at room temperature (RT) in paraformaldehyde (PFA; 4% w/v; Sigma) and sucrose (20% w/v; Sigma) solution in 1× PBS. The cells were then washed in PBS and incubated for 30 min at RT in goat serum (GS; 10% v/v; Invitrogen) in PBS to block non-specific staining. Neurons were then incubated for 60 min at RT with Flag antibody (mouse, 1:400, Sigma, cat #F3165) in PBS–GS blocking solution. After washing, neurons were incubated with Cy™3 AffiniPure Donkey Anti-Mouse IgG (H+L) (2.7 µg/ml; Jackson ImmunoResearch, cat #111-165-003) for standard epifluorescence assays in PBS-GS solution. The coverslips were then washed, and mounted on slides with mowiol 844 (48 mg/ml; Sigma).

##### Surface/total ratio for KCC2 or NKCC1

Cells were incubated for 30 minutes minimum at RT in normal goat serum (GS) (10%, v/v, Invitrogen) in PBS to block non-specific staining. KCC2-Flag or NKCC1-HA present at the neuronal surface were labeled by incubating cells for 1hour at RT with primary antibodies against Flag (mouse, 1:400, Sigma, cat #F3165) or HA (rabbit, 1:500, Cell signaling Technology, cat #C29F4) prepared in PBS-GS 3% solution. After three washes in PBS (1X), cells were incubated for 45 min at RT with CY3-conjugated donkey anti-mouse antibodies (2.7 µg/ml, 715-165-150, Jackson Immunoresearch, West Grove, USA) or anti-rabbit antibodies (2.5 µg/ml, 711-175-152, Jackson Immunoresearch, West Grove, USA) prepared in PBS-GS 3% solution to revel surface KCC2-Flag or NKCC1-HA. After three washes in PBS (1X), cells were permeabilized for 4 min at RT with Triton X-100 (0.25% w/v; Invitrogen) in PBS (1X). Cells were subsequently incubated for 1 h at RT with Flag or HA primary antibodies as above, in PBS-GS 3% solution to label total (surface + intracellular) KCC2-Flag or NKCC1-HA. After three washes in PBS (1X), cells were incubated for 45 min at RT with Alexa Fluor 647 AffiniPure Donkey Anti-mouse IgG (H + L) (2.7 µg/ml, Jackson ImmunoResearch, 715-605-150). or anti-rabbit antibodies (2.5 µg/ml, 711-165-152, Jackson Immunoresearch, West Grove, USA) to reveal total KCC2-Flag or NKCC1-HA. After three washes in PBS (1X), cells were finally mounted on glass slides using Mowiol 4–88 (48 mg/mL, Sigma).

#### Staining for colocalization experiments

##### For gephyrin and GABA_A_R γ2 colocalization

cells were washed 4 times with imaging medium at 4°C and then incubated for 20 minutes at 4°C with guinea pig GABA_A_R γ2 antibodies (1:500, cat #224 004, Synaptic Systems) in imaging medium. Cells were then fixed for 7 min with PFA-sucrose as described above, washed rapidly several times with PBS 1X and then permeabilized for 4 min at RT with Triton X-100 (0.25% w/v; Invitrogen) in PBS (1X). After several washes, cells were incubated for 30 min minimum at RT in normal goat serum (GS) (10%, v/v, Invitrogen) in PBS to block non-specific staining. Cells were incubated with mouse gephyrin antibodies (5 µg/ml, 147 011, Synaptic Systems) for 1h at room temperature. After three washes, cells were incubated for 45 min at RT with a mix of Alexa 488 conjugated donkey anti-mouse antibodies (1.6 µg/ml, 715-545-150, Jackson Immunoresearch) and CY3-conjugated goat anti-guineapig secondary antibodies (1.6 µg/ml, 106-165-003, Jackson Immunoresearch). After three washes, cells were finally mounted on glass slides using Mowiol 4–88 (48 mg/mL, Sigma).

##### For KCC2-Flag and NKCC1-HA colocalization

cells were fixed for 4 minutes with PFA-sucrose as described above, washed in PBS 1X and incubated for 30 minutes minimum at RT in normal GS (10%, v/v, Invitrogen) in PBS 1X. KCC2-Flag and NKCC1-HA present at the neuronal surface were labeled by incubating cells for 2 hours at RT with primary antibodies against Flag (mouse, 1:400, Sigma, cat #F3165) and HA (rabbit, 1:500, Cell signaling Technology, cat #C29F4) prepared in PBS-GS 3% solution. After three washes, neurons were incubated with Cy3 Donkey Anti-Rabbit IgG (H + L) (1:400; Jackson ImmunoResearch, France, 111-165-003) for standard epifluorescence assays and Alexa Fluor 647 AffiniPure Donkey Anti-mouse IgG (H + L) (1:400, Jackson ImmunoResearch, 715-605-150). After three washes, cells were finally mounted on glass slides using Mowiol 4–88 (48 mg/mL, Sigma).

#### Fluorescence image acquisition and analysis

Image acquisition was performed using a 63X objective (NA 1.32) on a Leica (Nussloch, Germany) DM6000 upright epifluorescence microscope with a 12-bit cooled CCD camera (Micromax, Roper Scientific, Evry, France) run by MetaMorph software (Roper Scientific). Image exposure time was determined on bright cells to obtain best fluorescence to noise ratio and to avoid pixel saturation. All images from a given culture were then acquired with the same exposure time and acquisition parameters.

##### For surface/total ratio for KCC2 or NKCC1

quantification was performed using the Meta Imaging 7.7 software (Molecular Devices, USA). A dendritic region of interest was manually chosen and the mean average intensity per pixel was measured.

##### For cluster colocalization analysis

quantification was performed using the Meta Imaging 7.7 software (Molecular Devices, USA). For each image, a dendritic region of interest was manually chosen. The images were then flattened, the background was filtered (kernel size, 3 × 3 × 2) to enhance cluster outlines, and a user-defined intensity threshold was applied to select clusters and avoid their coalescence.

For quantification of KCC2-Flag clusters colocalized with NKCC1-HA clusters, KCC2-Flag clusters comprising at least 2 pixels and colocalized on at least 1 pixel with NKCC1-HA clusters were considered. The number of clusters, the surface area and the integrated fluorescence intensities of clusters were measured for colocalized and non-colocalized clusters.

For quantification of GABA_A_R γ2 synaptic clusters, clusters comprising at least 2 pixels and colocalized on at least 1 pixel with gephyrin clusters were considered. The number of clusters, the surface area and the integrated fluorescence intensities of clusters were measured for colocalized and non-colocalized clusters.

Experimenters were blind to the condition of the sample analyzed. Sample size selection for experiments was based on published experiments, pilot studies as well as in-house expertise.

##### Chloride imaging

Neurons were imaged at 33 °C in a temperature-controlled open chamber (BadController V; Luigs & Neumann) mounted onto an Olympus IX71 inverted microscope equipped with a 60X objective (1.42 numerical aperture (NA); Olympus). CFP and YFP were detected using Lambda DG-4 monochromator (Sutter Instruments) coupled to the microscope through an optic fiber with appropriate filters (excitation, D436/10X and HQ485/15X; dichroic, 505DCXR; emission, HQ510lp; CFP and YFP filters from Chroma Technology). Images were acquired with an ImagEM EMCCD camera (Hamamatsu Photonics) and MetaFluor software (Roper Scientific). Mean background fluorescence (measured from a nonfluorescent area) was subtracted and the ratio F480/F440 was determined. Images (16-bit; 512 × 512) were typically acquired every 30 s for 5 minutes, with an integration time of 30-500 ms. Regions of interest (ROIs) were selected for measurement if they could only be measured over the whole experiment.

### Behavioral tests

Experimenters were blind to the condition of the groups analyzed. Group size selection for experiments was based on published experiments, pilot studies as well as in-house expertise. Two independent cohorts were used.

#### PTZ experiment

Pentylenetetrazole was administered as a single subcutaneous injection (60 mg/kg, volume <300 µL) to induce status epilepticus within a few minutes. Immediately after the injection, the animals were placed in a cage for one hour and their behavior was recorded on video. Behavioral manifestations of seizures were classified blindly according to the Racine scale (stage 0: no abnormal behavior; stage 1: immobility; stage 2: head nodding and rounded back; stage 3: lower limb clonic movements without rearing; stage 4: clonic rearing and uncontrolled jumping; stage 5: tonic-clonic seizure with loss of postural control and full extension of the fore and hind limbs (status epilepticus)). After one hour, the animals were euthanized.

#### Human postoperative epileptic tissues ex vivo experiments

Brain tissue specimens were obtained from the Neurosurgery center of GHU Paris Sainte-Anne hospital, Paris, France. All patients gave their written consent before the surgery. Surgery was not modified for the research protocol. All the brain tissue specimens came from the peritumoral tissue obtained from the security margin of glioma surgery. This study was approved by the Comité d’Evaluation et d’Ethique de l’INSERM - IRB00003888 (approval n°21-864). Tissues were prepared as already described ^70^. Planar MEA with titanium nitride electrodes (30 µm diameter, MultiChannel Systems) arranged in a 12 x 10 matrix with 1000 µm interelectrode spacing in the vertical axis, and 1500 µm in the horizontal axis were used to record extracellular activities. Recordings were made using a sampling rate of 10.000 Hz. Signals were filtered between 0.1 and 10.000 Hz (Logiciel MC Rack, MultiChannel Systems). To keep alive slices for several hours, the MEA chamber was perfused with a high flow (6 ml/min) of oxygenated aCSF at 36°C. Two different types of aCSF were used: during the first ten minutes, physiologic aCSF was used to record spontaneous activities (IIDs), and after ten minutes, ictogenic aCSF (124 mM NaCl, 6-8 mM KCl, 26 mM NaHCO3, 11.1 mM D-glucose, 1.6 mM CaCl2, 0.25 mM MgCl2) to elicit seizure-like events. Detection of seizure-like events and visualization of representative voltage traces were performed with SpikeSpector software (Cases-Cunillera et al., under review).

### Statistics

Unless otherwise stated, normally distributed data are presented as mean ± SEM (standard error of the mean), whereas non-normally distributed data are given as medians ± IQR (inter quartile range). For each experiment, statistical details (n, p-values, and test used) can always be found in the figure legends. All statistical analyses were performed using GraphPad Prism 10.1.0 (Dotmatics, Boston, USA). Normality of distribution was assessed by the Shapiro-Wilk test. Normally distributed unpaired datasets were compared using the unpaired t test, ordinary one-way ANOVA tests followed by Holm-Sidak’s multiple comparison tests or two-way ANOVA tests followed by Sidak’s multiple comparison for kinetic analyses. Non-Gaussian unpaired datasets were tested by two-tailed unpaired non-parametric Mann-Whitney test or Kruskal–Wallis tests followed by Dunn’s multiple comparison tests. Normally distributed paired datasets were compared using paired t-test, whereas non-Gaussian paired datasets were tested by the paired Wilcoxon test. Cumulative distributions were compared with the Kolmogorov-Smirnov test. Indications of significance corresponding to p-values < 0.05 (*), p < 0.01 (**), p < 0.001 (***) are reported in the figures and in the corresponding legends. Statistical analysis of MEA metrics was performed using Friedman’s test for repeated measures, followed by Conover post-hoc comparisons with Holm correction with the following significance levels: p < 0.05, *p < 0.01, **p < 0.001.

### Data and code availability

The data that support the findings of this study are available from the corresponding author upon reasonable request.

