## Supplemental Information for "NKCC1 as a signaling hub regulating KCC2 stability, chloride homeostasis, and seizure susceptibility"

### Supplementary Information

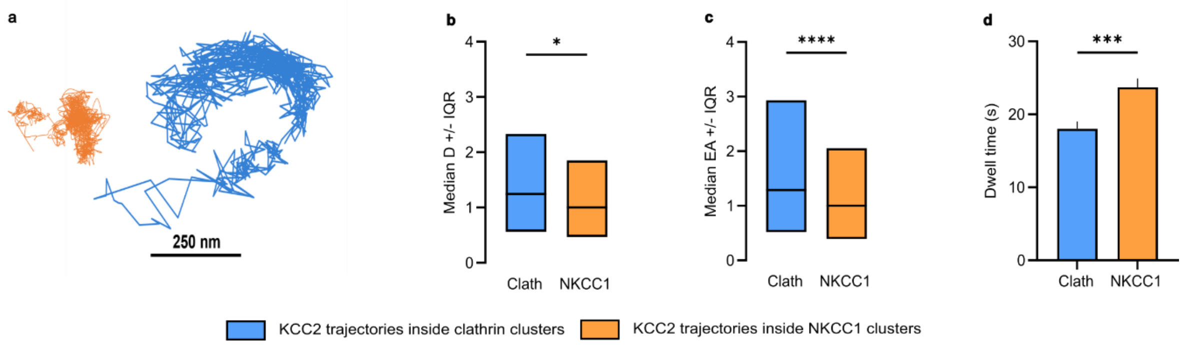


**Fig. S1. KCC2 are more slowed down and confined when they encounter NKCC1 clusters than clathrin clusters.**

**a.** Representative trajectories showing reduced surface exploration of KCC2 within NKCC1 clusters (orange), compared to KCC2 within endocytic zones labeled with clathrin-YFP (blue). Scale bar, 250 nm.

**b–c.** Median diffusion coefficient (D) (**b**) and explored area (EA) (**c**) of KCC2 within clathrin (blue) or NKCC1 (orange) clusters, shown as median ± interquartile range (25–75%). Values are normalized to the median KCC2 values within NKCC1 clusters.

**b.** Clathrin: $n = 310$ QDs; NKCC1: $n = 268$ QDs; 2 cultures. Kolmogorov–Smirnov (KS) test, $p = 0.0495$.

**c.** Clathrin: $n = 927$ QDs; NKCC1: $n = 804$ QDs; KS test, $p < 0.0001$.

**d.** Mean dwell time (DT) of KCC2 within clathrin (blue) or NKCC1 (orange) clusters (mean ± SEM).

Clathrin: $n = 106$ QDs; NKCC1: $n = 131$ QDs; Mann–Whitney (MW) test, $p = 0.0008$.

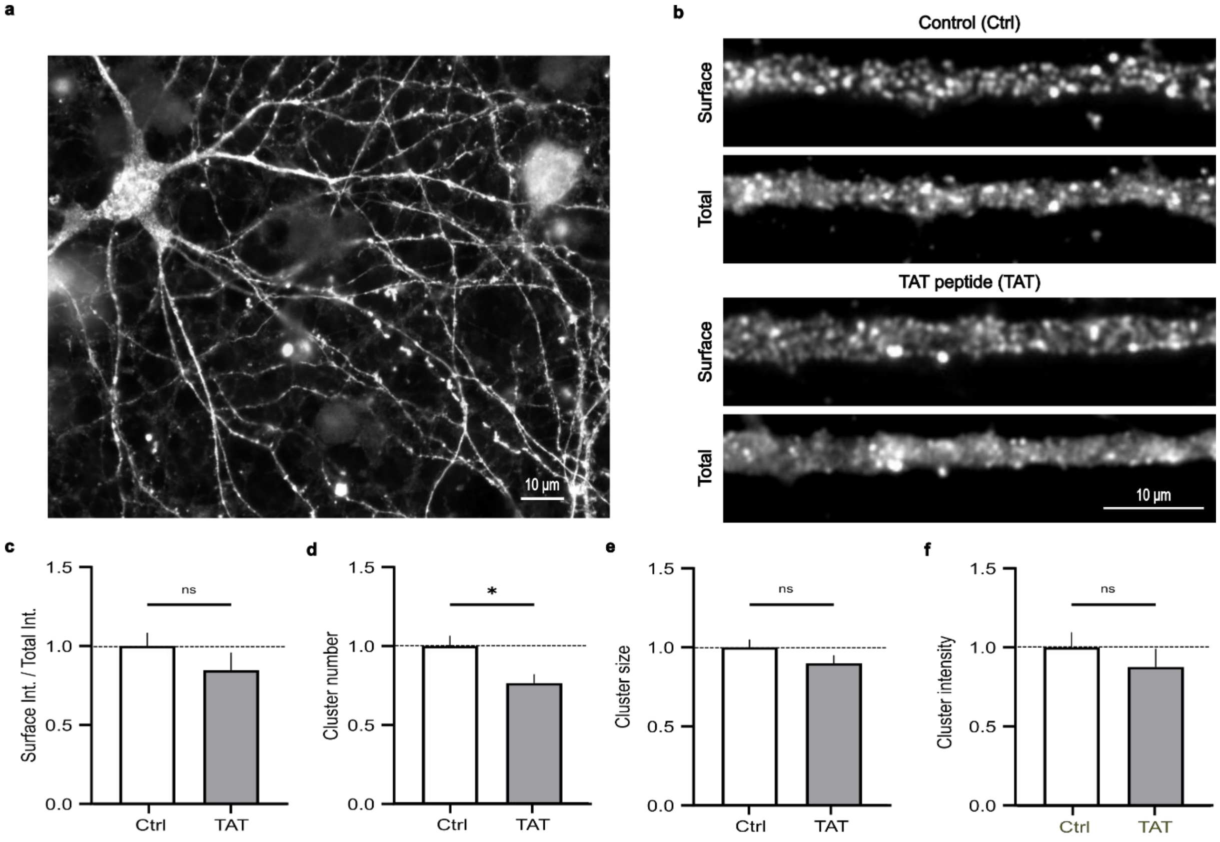


**Fig. S2. The cell-penetrating peptide TAT has a slight effect on KCC2 cluster number.**

**a.** Immunostaining of the biotinylated TAT peptide using fluorescent streptavidin in DIV21–23 hippocampal neurons after 1 hour incubation at 37°C. Scale bar, 10 μm.

**b.** Surface and total (surface + intracellular) Flag immunostaining of KCC2 in neurons expressing KCC2-Flag, in the absence or presence of TAT peptide (1 h, 100 μM). Scale bar, 10 μm.

**c.** Surface-to-total pixel intensity ratio of KCC2 remains unchanged in neurons treated (grey) or untreated (white) with TAT peptide.

Ctrl: $n$ = 40 cells; TAT: $n$ = 48 cells; 2 cultures. Mann–Whitney (MW) test: $p$ = 0.1033.

**d–f.** Number (**d**), size (**e**), and intensity (**f**) of KCC2 clusters in neurons treated with saline Ctrl: $n$ = 37 cells; TAT: $n$ = 37 cells; 2 cultures. MW test — cluster number: $p$ = 0.0239; cluster size: $p$ = 0.1883; cluster intensity: $p$ = 0.1608.

In all graphs (c–f), data are presented as mean ± SEM and normalized to their respective control values.

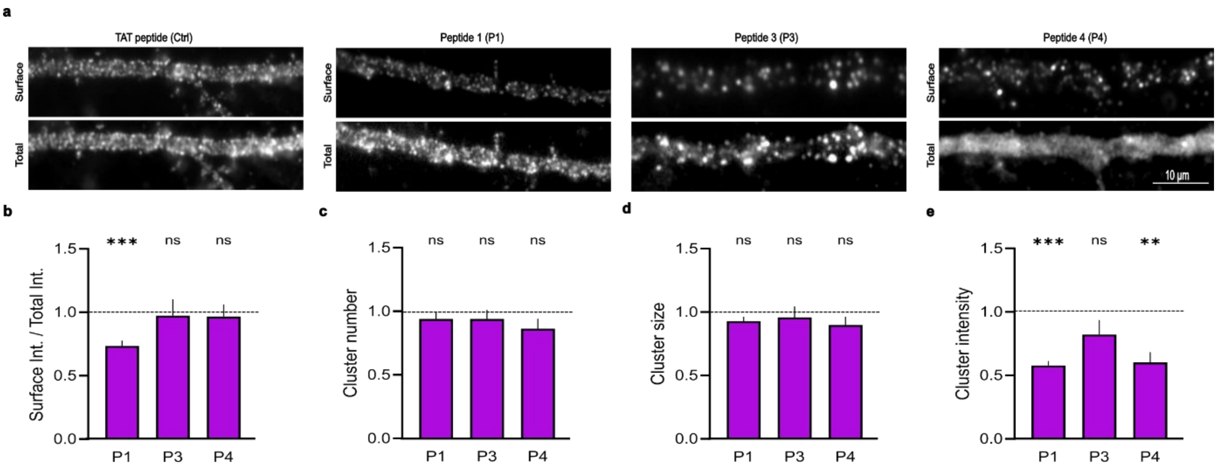


**Fig. S3. Peptides 1, 3, and 4 require the SPAK2-PP1 binding site for their activity.**

**a.** Surface and total (surface + intracellular) Flag immunostaining of KCC2 in neurons co-expressing KCC2 and NKCC1-ΔSPAK2ΔPP1 following 1-hour treatment with peptides TAT, P1, P3 or P4 (100 µM). Scale bar, 10 µm.

**b.** Surface-to-total pixel intensity ratio of KCC2 after peptide treatment.
P1: TAT, n = 81 cells; P1, n = 79 cells; 5 cultures; MW test: p = 0.0004.
P3: TAT, n = 35 cells; P3, n = 34 cells; 2 cultures; MW test: p = 0.4342.
P4: TAT, n = 35 cells; P4, n = 36 cells; 2 cultures; MW test: p = 0.8684.

**c–e.** Mean KCC2 cluster number (d), size (e), and intensity (f) in neurons co-expressing KCC2 and NKCC1-ΔSPAK2ΔPP1, following treatment with control peptide (TAT) or peptides P1, P3, or P4.

P1: TAT, n = 68 cells; P1, n = 64 cells; 3 cultures; MW test — cluster number: p = 0.2810; cluster size: p = 0.2433; cluster intensity: p < 0.0001.

P3: TAT, n = 28 cells; P3, n = 27 cells; 2 cultures; MW test — cluster number: p = 0.4568; cluster size: p = 0.5308; cluster intensity: p = 0.2952.

P4: TAT, n = 28 cells; P4, n = 27 cells; 2 cultures; MW test — cluster number: p = 0.1083; cluster size: p = 0.2441; cluster intensity: p = 0.0081.

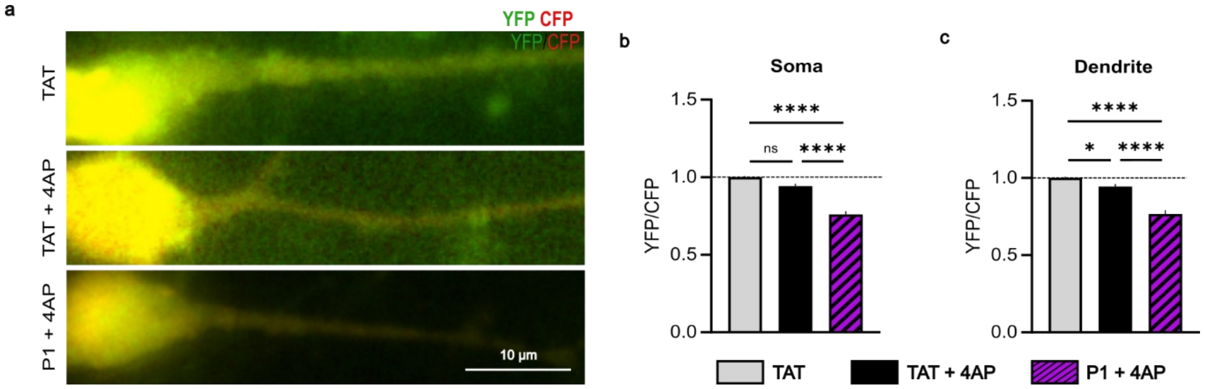


**Fig. S4. Peptide 1 potentiates 4AP-induced elevation of intracellular chloride levels.**

**a.** Representative overlay images of CFP (red) and YFP (green) in neurons expressing SuperClomeleon, NKCC1, and KCC2 under Control TAT, TAT + 4AP, and P1 + 4AP conditions. Scale bar, 10 µm.

**b-c.** YFP/CFP fluorescence ratios measured in the soma (b) and dendrite (c) of neurons expressing SuperClomeleon, NKCC1, and KCC2 under Control (TAT, grey), TAT + 4AP (black), and P1 + 4AP (hatched pink) conditions.

TAT, n = 39 cells; TAT + 4AP, n = 42 cells; P1 + 4AP, n = 23 cells; 3 cultures;

One-way ANOVA test:

Soma: TAT vs TAT+4AP $p = 0.0760$; TAT+4AP vs P1 + 4AP $p < 0.0001$; TAT vs P1 + 4AP $p < 0.0001$.

Dendrite: TAT vs TAT+4AP $p = 0.0323$; TAT+4AP vs P1 + 4AP $p < 0.0001$; TAT vs P1 + 4AP $p < 0.0001$.

All data are presented as mean ± SEM. Values were normalized to the respective TAT control condition.

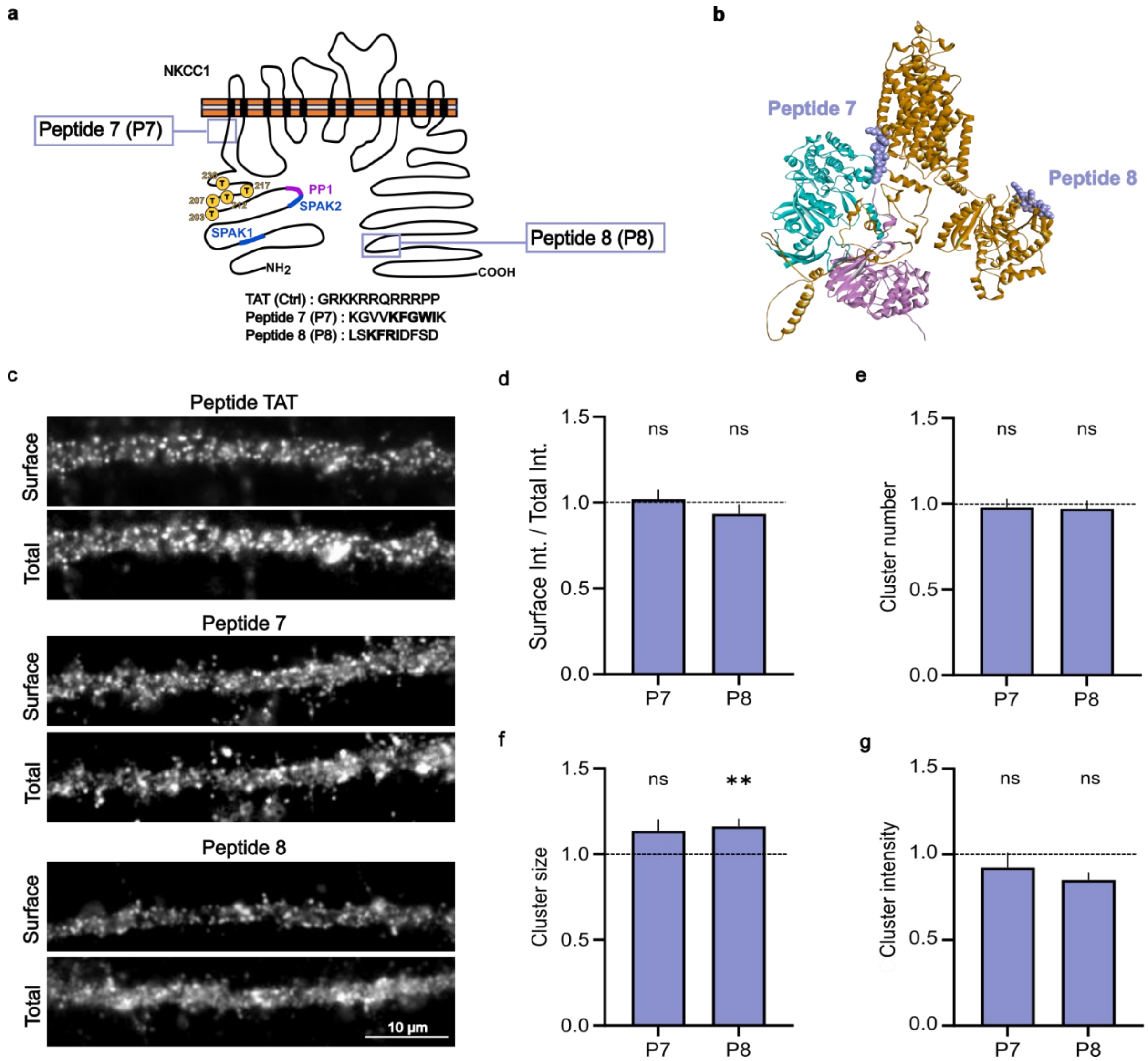


**Fig. S5. No effect of peptides 7 and 8, which mimic NKCC1 regions containing putative SPAK recognition sequences distant from the SPAK1 and SPAK2 binding sites.**

**a.** Schematic representation of NKCC1 indicating the positions and amino acid sequences of peptides 7 (P7) and 8 (P8).

**b.** Structure of NKCC1 (brown), SPAK (blue), and PP1 (pink). The grey regions correspond to peptide sequences 7 and 8.

**c.** Surface and total (surface + intracellular) Flag immunostaining of KCC2 in neurons co-expressing NKCC1 and KCC2, after 1-hour treatment with control peptide (TAT) or peptides P7 or P8 (100 µM). Scale bar, 10 µm.

**d.** Surface-to-total pixel intensity ratio of KCC2 after treatment with peptides P7 or P8, compared to the TAT control condition.

P7: TAT, $n$ = 27 cells; P7, $n$ = 22 cells; 2 cultures. MW test: $p$ = 0.7880.

P8: TAT, $n$ = 27 cells; P8, $n$ = 22 cells; 2 cultures. MW test: $p$ = 0.5974.

**e–g.** Mean number (**d**), size (**e**), and intensity (**f**) of KCC2 clusters in neurons treated with P7 or P8, compared to the TAT control.

Data are shown as mean ± SEM and normalized to their respective control TAT values.

P7: TAT, *n* = 27 cells; P7, *n* = 22 cells (ok chiffres vérifiés); 2 cultures. MW test — cluster number: *p* = 0.6396; cluster size: *p* = 0.1200; cluster intensity: *p* = 0.3328.

P8: TAT, *n* = 40 cells; P8, *n* = 28 cells; 3 cultures. MW test — cluster number: *p* = 0.7336; cluster size: *p* = 0.0035; cluster intensity: *p* = 0.0989.

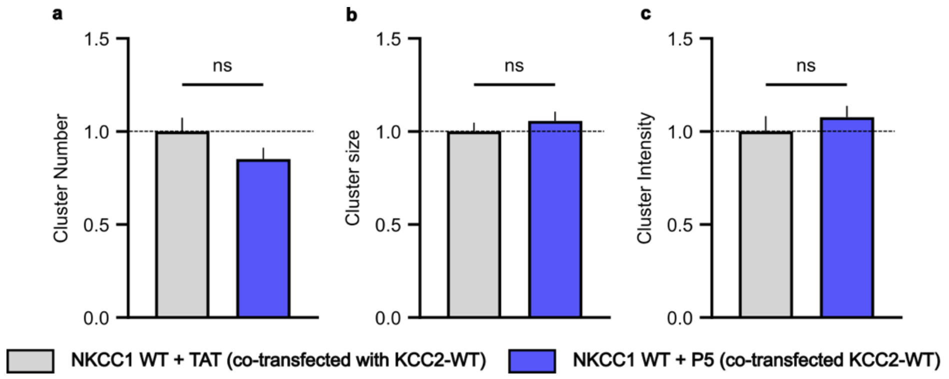


**Fig. S6. Peptide 5 has no effect on NKCC1 clustering.**

Mean NKCC1 cluster number (**a**), size (**b**), and intensity (**c**) in neurons expressing NKCC1-WT and KCC2-WT, treated with peptides TAT or P5.

NKCC1-WT: TAT, n = 23 cells; P5, n = 27 cells; 2 cultures; MW test — cluster number: p = 0.1566; cluster size: p = 0.5334; cluster intensity: p = 0.2698.

All data are presented as mean ± SEM. Values were normalized to their respective control (TAT) condition.

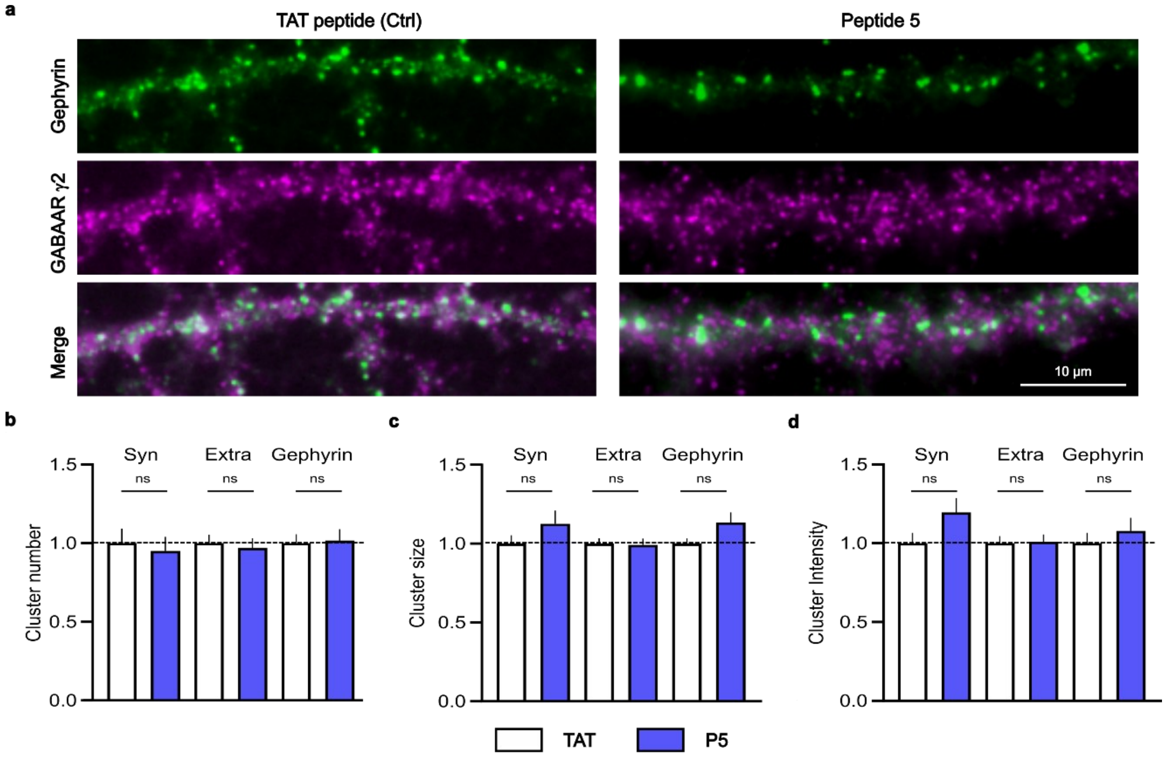


**Fig. S7. Peptide 5 has no effect on $GABA_AR$ and gephyrin clustering.**

**a.** Immunostaining of the $GABA_A$ receptor γ2 subunit (purple) and gephyrin (green) in DIV21 hippocampal neurons treated with peptide 5 (P5), compared to the control condition (TAT). Scale bar: 10 µm.

**b–d.** Quantification of the mean number (b), size (c), and intensity (d) of synaptic (Syn) and extrasynaptic (Extra) $GABA_A$ receptor γ2 clusters colocalized or not with gephyrin, as well as all gephyrin clusters, in neurons treated with P5 (blue) or control TAT peptide (white).

Data are shown as mean ± SEM and normalized to their respective control values.

TAT: n = 35 cells; P5: n = 34 cells; 3 cultures.

Synaptic $GABA_A$ receptor γ2: Mann–Whitney (MW) test — cluster number: $p = 0.7429$; cluster size: $p = 0.4795$; cluster intensity: $p = 0.0850$.

Extrasynaptic $GABA_A$ receptor γ2: MW test — cluster number: $p = 0.5232$; cluster size: $p = 0.6121$; cluster intensity: $p = 0.8627$.

Gephyrin: MW test — cluster number: $p = 0.2388$; cluster size: $p < 0.0001$; cluster intensity: $p = 0.6633$.

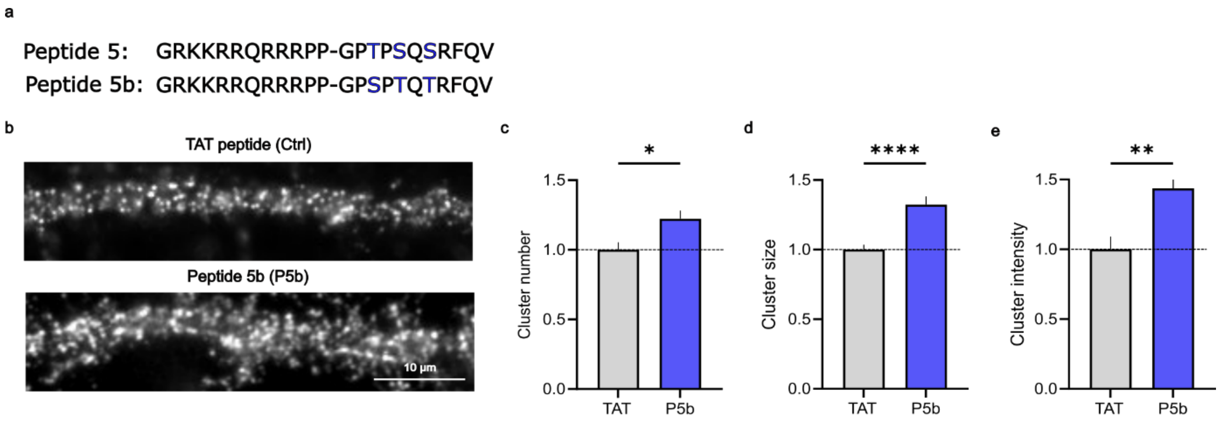


**Fig. S8. Peptides 5b increases KCC2 clustering.**

**a.** Amino acid sequences of peptides 5 (P5) and 5b (P5b).

**b.** KCC2 surface expression after treatment with peptide 5b P5b (1 hour, 100 µM), compared to the control TAT peptide. Scale bar, 10 µm.

**c–e.** Peptide P5b increases the mean number (**c**), size (**d**), and intensity (**e**) of KCC2 clusters compared to the TAT control peptide.

Data are presented as mean ± SEM and normalized to their respective TAT control values.

TAT, $n$ = 31 cells; P5b, $n$ = 38 cells; 3 cultures.

MW test — cluster number: $p = 0.0130$; cluster size: $p < 0.0001$; cluster intensity: $p = 0.0028$.
